# The genome of *Ectocarpus subulatus* – a highly stress-tolerant brown alga

**DOI:** 10.1101/307165

**Authors:** Simon M. Dittami, Erwan Corre, Loraine Brillet-Guéguen, Agnieszka P. Lipinska, Noé Pontoizeau, Meziane Aite, Komlan Avia, Christophe Caron, Chung Hyun Cho, Jonas Collén, Alexandre Cormier, Ludovic Delage, Sylvie Doubleau, Clémence Frioux, Angélique Gobet, Irene González-Navarrete, Agnès Groisillier, Cécile Hervé, Didier Jollivet, Hetty KleinJan, Catherine Leblanc, Xi Liu, Dominique Marie, Gabriel V. Markov, André E. Minoche, Misharl Monsoor, Pierre Pericard, Marie-Mathilde Perrineau, Akira F. Peters, Anne Siegel, Amandine Siméon, Camille Trottier, Hwan Su Yoon, Heinz Himmelbauer, Catherine Boyen, Thierry Tonon

**Author notes:** Deceased (Christophe Caron).

## Abstract

Brown algae are multicellular photosynthetic stramenopiles that colonize marine rocky shores worldwide. *Ectocarpus* sp. Ec32 has been established as a genomic model for brown algae. Here we present the genome and metabolic network of the closely related species, *Ectocarpus subulatus* Kützing, which is characterized by high abiotic stress tolerance. Since their separation, both strains show new traces of viral sequences and the activity of large retrotransposons, which may also be related to the expansion of a family of chlorophyll-binding proteins. Further features suspected to contribute to stress tolerance include an expanded family of heat shock proteins, the reduction of genes involved in the production of halogenated defence compounds, and the presence of fewer cell wall polysaccharide-modifying enzymes. Overall, *E. subulatus* has mainly lost members of gene families down-regulated in low salinities, and conserved those that were up-regulated in the same condition. However, 96% of genes that differed between the two examined *Ectocarpus* species, as well as all genes under positive selection, were found to encode proteins of unknown function. This underlines the uniqueness of brown algal stress tolerance mechanisms as well as the significance of establishing *E. subulatus* as a comparative model for future functional studies.

## Introduction

Brown algae (Phaeophyceae) are multicellular photosynthetic organisms that are successful colonizers of rocky shores in the world’s oceans. In many places they constitute the dominant vegetation in the intertidal zone, where they have adapted to multiple stressors including strong variations in temperature, salinity, irradiation, and mechanical stress (wave action) over the tidal cycle^1^. In the subtidal environment, brown algae form kelp forests that harbor highly diverse communities^2^. They are also harvested as food or for industrial purposes, such as the extraction of alginates^3^. The worldwide annual harvest of brown algae has reached 10 million tons in 2014 and is constantly growing^4^. Brown algae share some basic photosynthetic machinery with land plants, but their plastids derived from a secondary or tertiary endosymbiosis event with a red alga, and they belong to an independent lineage of eukaryotes, the stramenopiles^5^. This phylogenetic background, together with their distinct habitat, contributes to the fact that brown algae have evolved numerous unique metabolic pathways, life cycle features, and stress tolerance mechanisms.

To enable functional studies of brown algae, strain Ec32 of the small filamentous alga *Ectocarpus* sp. has been established as a genetic and genomic model^6–8^. This strain was formerly described as *Ectocarpus siliculosus*, but has since been shown to belong to an independent clade by molecular methods^9,10^. More recently, three additional brown algal genomes, that of the kelp species *Saccharina japonica*^11^, that of *Cladosiphon okamuranus*^12^, and that of *Nemacystus decipiens*^13^, have been characterized. Comparisons between these four genomes have allowed researchers to obtain a first overview of the unique genomic features of brown algae, as well as a glimpse of the genetic diversity within this group. However, given the evolutionary distance between these algae, it is difficult to link genomic differences to physiological differences and possible adaptations to their lifestyle. To be able to generate more accurate hypotheses on the role of particular genes and genomic features for adaptive traits, a common strategy is to compare closely related strains and species that differ only in a few genomic features. The genus *Ectocarpus* is particularly well suited for such comparative studies because it comprises a wide range of morphologically similar but genetically distinct strains and species that have adapted to different marine and brackish water environments^9,14–16^. One species within this group, *Ectocarpus subulatus* Kützing^10^, comprises isolates highly resistant to elevated temperature^17^ and low salinity. A strain of this species was even isolated from freshwater^18^, constituting one of the handful of known marine-freshwater transitions in brown algae^19^.

Here we present the draft genome and metabolic network of a strain of *E. subulatus*, establishing the genomic basis for its use as a comparative model to study stress tolerance mechanisms, and in particular low salinity tolerance, in brown algae. Similar strategies have been successfully employed in terrestrial plants, where “extremophile” relatives of model- or economically relevant species have been sequenced to explore new stress tolerance mechanisms in the green lineage^20–25^. The study of the *E. subulatus* genome, and subsequent comparative analysis with other brown algal genomes, in particular that of *Ectocarpus* sp. Ec32, provides insights into the dynamics of *Ectocarpus* genome evolution and divergence, and highlights important adaptive processes, such as a potentially retrotransposon driven expansion of the family of chlorophyll-binding proteins with subsequent diversification. Most importantly, our analyses underline that most of the observed differences between the examined species of *Ectocarpus* correspond to proteins with yet unknown functions.

## Results

### Sequencing and assembly of the *E. subulatus* genome

A total of 34.7Gb of paired-end read data and of 28.8Gb of mate-pair reads (corresponding to 45 million non-redundant mate-pairs) were acquired (Supporting Information Table S1). The final genome assembly size of strain Bft15b was 227Mb (Table 1), and we also obtained 123Mb of bacterial contigs corresponding predominantly to *Alphaproteobacteria* (50%, with the dominant genera *Roseobacter* 8% and *Hyphomonas* 5%), followed by *Gammaproteobacteria* (18%), and *Flavobacteria* (13%). The mean sequencing coverage of mapped reads was 67X for the paired-end library, and the genomic coverage was 6.9, 14.4, and 30.4X for the 3kb, 5kb, and 10kb mate-pair libraries, respectively. RNA-seq experiments yielded 8.8Gb of RNA-seq data, of which 96.6% (Bft15b strain in seawater), 87.6% (freshwater strain in seawater), and 85.3% (freshwater strain in freshwater) aligned with the final genome assembly of the Bft15b strain.

**Table 1:** Assembly statistics of available brown algal genomes. PE = paired-end, MP = mate-pair, n.d. = not determined

|  | *E. subulatus*<br>Bft15b | *Ectocarpus*<br>sp. Ec32<sup>7</sup> | *S. japonica*<sup>11</sup> | *C. okamuranus*<sup>12</sup> | *N. decipiens*<sup>13</sup> |
| --- | --- | --- | --- | --- | --- |
| Sequencing strategy | Illumina<br>(PE+MP) | Sanger+Bac<br>libraries | Illumina<br>PE+PacBio | Illumina<br>(PE+MP) | Illumina<br>(PE+MP) |
| Genome size estimate<br>(flow cytometry) | 226 | 214<sup>6*</sup> | 545 | 140 | n. d. |
| Genome size<br>(assembled) | 242 Mb | 196 Mb | 537 Mb | 130 Mb | 154 Mb |
| Genomic Coverage | 119 X | 11 X<sup>#</sup> | 178 X | 100 X | 420 X |
| G/C contents | 54% | 53% | 50% | 54% | 56% |
| Number of scaffolds<br>>2kb | 1,757 | 1,561 | 6,985 | 541 | 685 |
| Scaffold N50 (kb) | 510 kb | 497 kb | 254 kb | 416 kb | 1,863 kb |
| Number of predicted<br>genes | 25,893 | 17,418 | 18,733 | 13,640 | 15,156 |
| Mean number of exons<br>per gene | 5.4 | 8.0 | 6.5 | 9.3 | 11.2 |
| Repetitive elements | 30% | 30%<sup>##</sup> | 40% | 4.1% | 8.8% |
| BUSCO genome<br>completeness<br>(complete+fragmented) | 86%<br>(91%<sup>*#</sup>) | 94%<br>(99%<sup>*#</sup>) | 91%<br>(96%<sup>*#</sup>) | 88% (93%<sup>*#</sup>) | 92%<br>(97%<sup>*#</sup>) |
| BUSCO Fragmented<br>proteins | 13.5% | 7.4% | 14.2% | 11.9% | 5.6% |

<sup>##</sup> 23% according to <sup>7</sup>, but 30% when re-run with the current version (2.5) of the REPET pipeline.
<sup>\*#</sup> not considering proteins absent from all three brown algal genomes.

### Gene prediction and annotation

The number of predicted proteins in *E. subulatus* was 60% higher than that predicted for Ec32 (Table 1), mainly due to the presence of mono-exonic genes, many of which corresponded to transposases, which were not removed from our predictions, but had been manually removed from the Ec32 genome. Ninety-eight percent of the gene models were supported by at least one associated RNA-seq read, and 92% were supported by at least ten reads, with lowly-expressed (<10 reads) genes being generally shorter (882 vs 1,403 bases), and containing fewer introns (2.6 vs 5.7). In 7.3% of all predicted proteins we detected a signal peptide, and 3.7% additionally contained an ‘ASAFAP’-motif (Supporting Information Table S2) indicating that they are likely targeted to the plastid^26^. Overall the BUSCO^27^ analyses indicate that the *E. subulatus* genome is 86% complete (complete and fragmented genes) and 91% when not considering proteins also absent from all other currently sequenced brown algae (Table 1).

### Repeated elements

Thirty percent of the E. subulatus genome consisted of repeated elements. The most abundant groups of repeated elements were large retrotransposon derivatives (LARDs), followed by long terminal repeats (LTRs, predominantly Copia and Gypsy), and long and short interspersed nuclear elements (LINEs, Figure 1A). The overall distribution of sequence identity levels within superfamilies showed two peaks, one at an identity level of 78-80%, and one at 96-100% (Figure 1C). An examination of transposon conservation at the level of individual families revealed a few families that follow this global bimodal distribution (*e.g.* TIR B343 or LARD B204), while the majority exhibited a unimodal distribution with peaks either at high (*e.g.* LINE R15) or at lower identity levels (*e.g.* LARD B554) (Figure 1C). Terminal repeat retrotransposons in miniature (TRIM) and LARDs, both non-autonomous groups of retrotransposons, were among the most conserved families. A detailed list of transposons is provided in Supporting Information Table S3. In line with previous observations carried out in *Ectocarpus* sp. Ec32, no methylation was detected in the *E. subulatus* genomic DNA.

**Figure 1:**
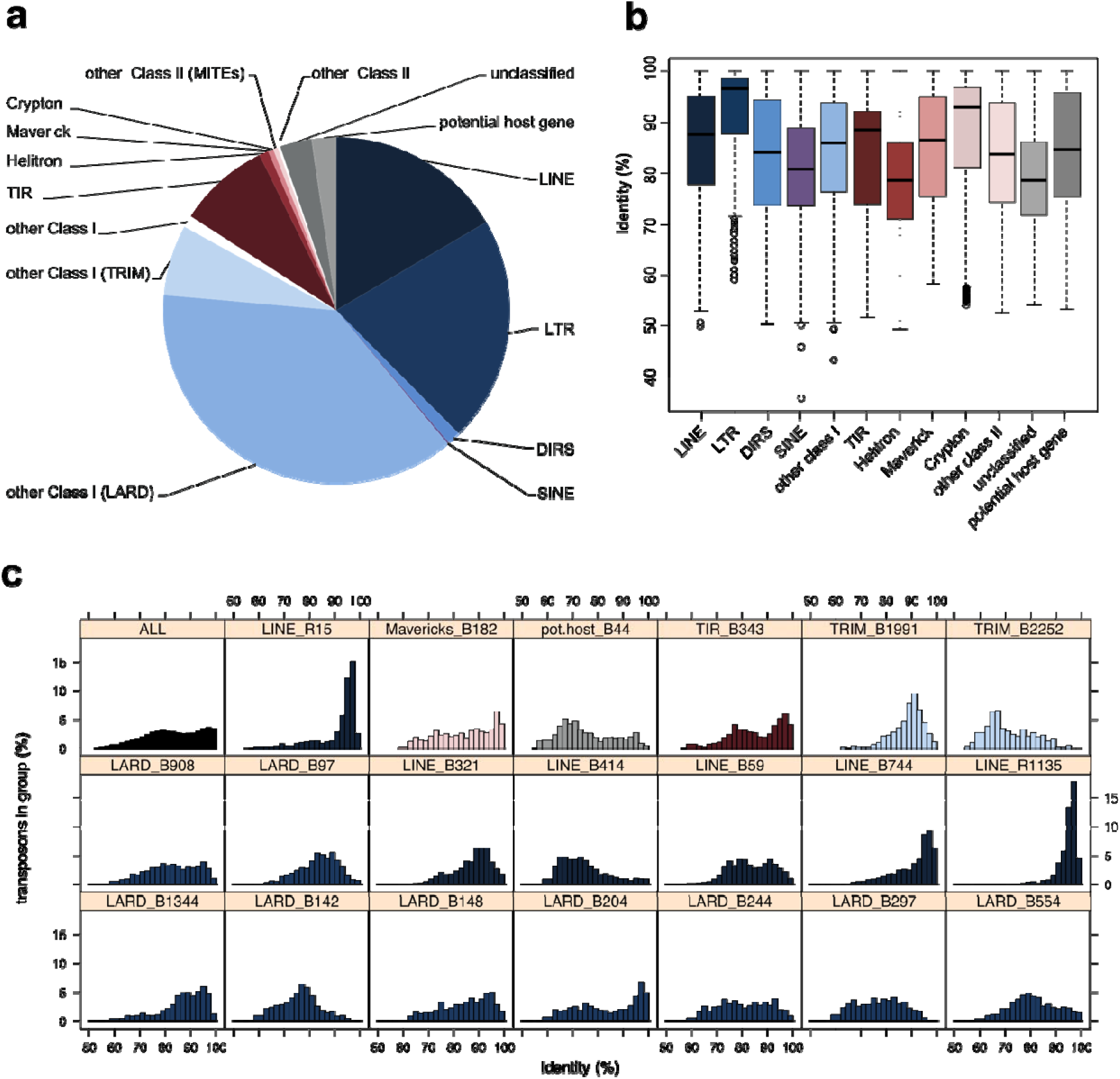
Repeated elements identified within the genome of *E. subulatus* Bft15b. A) Number of transposons detected in the different superfamilies; B) Boxplot of sequence identity levels for the detected superfamilies; and C) Distribution of sequence identities in all and the 20 most abundant transposon families.

### Organellar genomes

Plastid and mitochondrial genomes from *E. subulatus* have 95.5% and 91.5% sequence identity with their *Ectocarpus* sp. Ec32 counterparts in the conserved regions respectively. Only minor structural differences were observed between organellar genomes of both *Ectocarpus* genomes, as detailed in Supporting Information Text S1.

### Global comparison of predicted proteomes

#### Metabolic network-based comparisons

Similar to the network previously obtained for *Ectocarpus* sp. Ec32^28^, the *E. subulatus* Bft15b metabolic network comprised 2,074 metabolic reactions and 2,173 metabolites in 464 pathways, which can be browsed at http://gem-aureme.irisa.fr/sububftgem. In total, 2,445 genes associated with at least one metabolic reaction, and 215 pathways were complete (Figure 2). Comparisons between both networks were carried out on a pathway level (Supporting Information Text S1, Section “Metabolic network-based comparisons”), but no pathways were found to be truly specific to either Ec32 and/or Bf15b.

**Figure 2:**
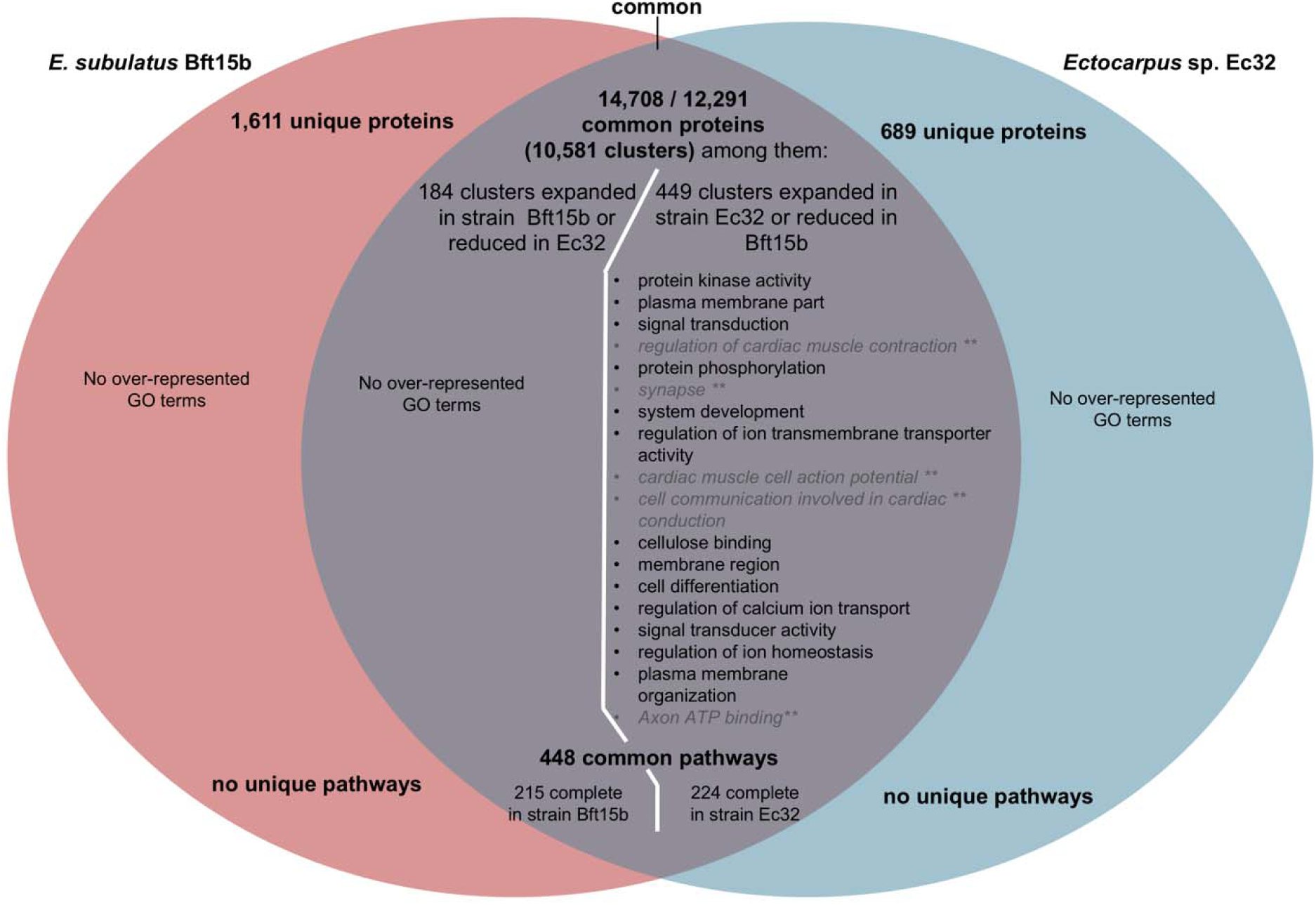
Comparison of gene content and metabolic capacities of *E. subulatus* Bft15b and *Ectocarpus* sp. Ec32. The top part of the Venn diagram displays the number of predicted proteins and protein clusters unique and common to both genomes in the OrthoFinder analysis. The middle part shows GO annotations significantly enriched (FDR ≤ 0.05) among these proteins. For the common clusters, the diagram also contains the results of gene set enrichment analyses for annotations found among clusters expanded in *E. subulatus* Bft15b and those expanded in *Ectocarpus* sp. Ec32. Functional annotations not directly relevant to the functioning of *Ectocarpus* or shown to be false positives are shown in grey and italics. The bottom part shows the comparison of both genomes in terms of their metabolic pathways.

#### Genes under positive selection

Out of the 2,311 orthogroups with single-copy orthologs that produced high quality alignments, 172 gene pairs (7.4%) exhibited dN/dS ratios > 0.5 (Supporting Information Table S4). Among these, only eleven (6.4%) were found to fit significantly better with the model allowing for positive selection in the *Ectocarpus* branch. These genes are likely to have been under positive selection, and two of them contained a signal peptide targeting the plastid. All of them are genes specific to the brown algal lineage with unknown function, and only two genes contained protein domains related to a biochemical function (one oxidoreductase-like domain, and one protein prenyltransferase, alpha subunit). However, all of them were expressed at least in *E. subulatus* Bft15b. There was no trend for these genes to be located in specific regions of the genome (all except two for *Ectocarpus* sp. Ec32 were on different scaffolds) and none of the genes were located in the pseudoautosomal region of the sex chromosome.

#### Genes specific to either *Ectocarpus* genome, and expanded genes and gene families

After manual curation based on tblastn searches to eliminate artefacts arising from differences in the gene predictions, 184 expanded gene clusters and 1,611 predicted proteins were found to be specific to *E. subulatus* compared to *Ectocarpus* sp., while 449 clusters were expanded and 689 proteins were found specifically in the latter (Figure 2, Supporting Information Table S5). This is far less than the 2,878 and 1,093 unique clusters found for a recent comparison of *N. decipiens* and *C. okamuranus*^13^. Gene set enrichment analyses revealed no GO categories to be significantly over-represented among the genes unique to or expanded in *E. subulatus* Bft15b, but several categories were over-represented among the genes and gene families specific to or expanded in the *Ectocarpus* sp. Ec32 strain. Many were related either to signalling pathways or to the membrane and transporters (Figure 2), but it is difficult to distinguish between the effects of a potentially incomplete genome assembly and true gene losses in Bft15b. In the manual analyses we therefore focussed on the genes specific to and expanded in *E. subulatus*.

Among the 1,611 *E. subulatus*-specific genes, 1,436 genes had no homologs (e-value < 1e-5) in the UniProt database as of May 20^th^ 2016: they could thus, at this point in time, be considered lineage-specific and had no function associated to them. Among the remaining 175 genes, 145 had hits (e-value < 1e-5) in *Ectocarpus* sp. Ec32, *i.e*. they likely correspond to multi-copy genes that had diverged prior to the separation of *Ectocarpus* and *S. japonica*, and for which the *Ectocarpus* sp. Ec32 and *S. japonica* orthologs were lost. Thirteen genes had homology only with uncharacterized proteins or were too dissimilar from characterized proteins to deduce hypothetical functions; another eight probably corresponded to short viral sequences integrated into the algal genome (EsuBft1730_2, EsuBft4066_3, EsuBft4066_2, EsuBft284_15, EsuBft43_11, EsuBft551_12, EsuBft1883_2, EsuBft4066_4), and one (EsuBft543_9) was related to a retrotransposon. Two adjacent genes (EsuBft1157_4, EsuBft1157_5) were also found in diatoms and may be related to the degradation of cellobiose and the transport of the corresponding sugars. Two genes, EsuBft1440_3 and EsuBft1337_8, contained conserved motifs (IPR023307 and SSF56973) typically found in toxin families. Two more (EsuBft1006_6 and EsuBft308_11) exhibited low similarities to animal and fungal transcription factors, and the last (EsuBft36_20 and EsuBft440_20) consisted almost exclusively of short repeated sequences of unknown function (“ALEW” and “GAAASGVAGGAVVVNG”, respectively). In total, 1.7% contained a signal peptide targeting the plastid, *i.e.* significantly less than the 3.7% in the entire dataset (Fisher exact test, p<0.0001).

The large majority of *Ectocarpus* sp. Ec32-specific proteins (511) also corresponded to proteins of unknown function without matches in public databases. Ninety-seven proteins were part of the *E. siliculosus* virus-1 (EsV-1) inserted into the Ec32 genome and the remaining 81 proteins were poorly annotated, usually only via the presence of a domain. Examples are ankyrin repeat-containing domain proteins (12), Zinc finger domain proteins (6), proteins containing wall sensing component (WSC) domains (3), protein kinase-like proteins (3), and Notch domain proteins (2).

Regarding the 184 clusters of expanded genes in *E. subulatus,* 139 (1,064 proteins) corresponded to proteins with unknown function, 98% of which were found only in *Ectocarpus*. Furthermore, nine clusters (202 proteins) represented sequences related to transposons predicted in both genomes, and eight clusters (31 proteins) were similar to known viral sequences. Only 28 clusters (135 proteins) could be roughly assigned to biological functions (Table 2). They comprised proteins potentially involved in modification of the cell-wall structure (including sulfation), in transcriptional regulation and translation, in cell-cell communication and signalling, as well as a few stress response proteins, notably a set of HSP20s, and several proteins of the light-harvesting complex (LHC) potentially involved in non-photochemical quenching. Only 0.6% of all genes expanded in Bft15b contained a signal peptide targeting the plastid, i.e. significantly less than the 3.7% in the entire dataset (Fisher exact test, p<0.0001).

**Table 2:** Clusters of orthologous genes identified by OrthoFinder as expanded in the genome of *E. subulatus* Bft15b or reduced in *Ectocarpus* sp. Ec32, after manual identification of false positives, and removal of clusters without functional annotation or related to transposon or viral sequences.

| Cluster(s) | # Ec32 | # Bft15b | Putative annotation or functional domain |
| --- | --- | --- | --- |
| *Cell-wall related proteins* |  |  |  |
| OG0000597 | 1 | 3 | Peptidoglycan-binding domain |
| OG0000284, -782, -118 | 6 | 12 | Carbohydrate-binding WSC domain |
| OG0000889 | 1 | 2 | Cysteine desulfuration protein |
| OG0000431 | 1 | 3 | Galactose-3-O-sulfotransferase (partial) |
| *Transcriptional regulation and translation* |  |  |  |
| OG0000785 | 1 | 2 | AN1-type zinc finger protein |
| OG0000059 | 4 | 10 | C2H2 zinc finger protein |
| OG0000884 | 1 | 2 | Zinc finger domain |
| OG0000766 | 1 | 2 | DNA-binding SAP domain |
| OG0000853 | 1 | 2 | RNA binding motif protein |
| OG0000171 | 1 | 6 | Helicase |
| OG0000819 | 1 | 2 | Fungal transcriptional regulatory protein domain |
| OG0000723 | 1 | 2 | Translation initiation factor eIF2B |
| OG0000364 | 2 | 3 | Ribosomal protein S15 |
| OG0000834 | 1 | 2 | Ribosomal protein S13 |
| *Cell-cell communication and signaling* |  |  |  |
| OG0000967 | 1 | 2 | Ankyrin repeat-containing domain |
| OG0000357 | 2 | 3 | Regulator of G protein signaling domain |
| OG0000335 | 2 | 3 | Serine/threonine kinase domain |
| OG0000291 | 2 | 3 | Protein kinase |
| OG0000185 | 3 | 4 | Octicosapeptide/Phox/Bem1p domain |
| *Others* |  |  |  |
| OG0000726 | 1 | 3 | HSP20 |
| OG0000104 | 1 | 9 | Light harvesting complex protein |
| OG0000277 | 3 | 3 | Major facilitator superfamily transporter |
| OG0000210 | 2 | 4 | Cyclin-like domain |
| OG0000721 | 1 | 2 | Myo-inositol 2-dehydrogenase |
| OG0000703 | 1 | 2 | Short-chain dehydrogenase |
| OG0000749 | 1 | 2 | Putative Immunophilin |
| OG0000463 | 1 | 3 | Zinc-dependent metalloprotease with notch domain |

Striking examples of likely expansions in *Ectocarpus* sp. Ec32 or reduction in *E. subulatus* Bft15b were different families of serine-threonine protein kinase domain proteins present in 16 to 25 copies in Ec32 compared to only 5 or 6 in Bft15b, Kinesin light chain-like proteins (34 vs. 13 copies), two clusters of Notch region containing proteins (11 and 8 vs. 2 and 1 copies), a family of unknown WSC domain containing proteins (8 copies vs. 1), putative regulators of G-protein signalling (11 vs. 4 copies), as well as several expanded clusters of unknown and viral proteins. However, these results need to be taken with caution because the *E. subulatus* Bft15b genome was less complete than that of *Ectocarpus* sp. Ec32.

#### Correlation with gene expression patterns

To assess whether genomic adaptations in *E. subulatus* Bft15b were located preferentially in genes that are known to be responsive to salinity stress, we compared expanded gene families to previously available expression data obtained for a freshwater strain of *E. subulatus* grown in freshwater vs seawater^29^. This analysis revealed that genes that were down-regulated in response to low salinity were significantly over-represented among the gene families expanded in *Ectocarpus* sp. Ec32 or reduced in *E. subulatus* Bft15b, (42% of genes vs 26% for all genes; Fischer exact test p=0.0002), while genes that were upregulated in response to low salinity were significantly under-represented (25% vs 33%; Fischer exact test p=0.006; Figure 3, Supporting Information Table S6). This indicates that *E. subulatus* Bft15b has mainly lost members of gene families that were generally down-regulated in low salinities, and conserved those that were upregulated in this condition.

**Figure 3:**
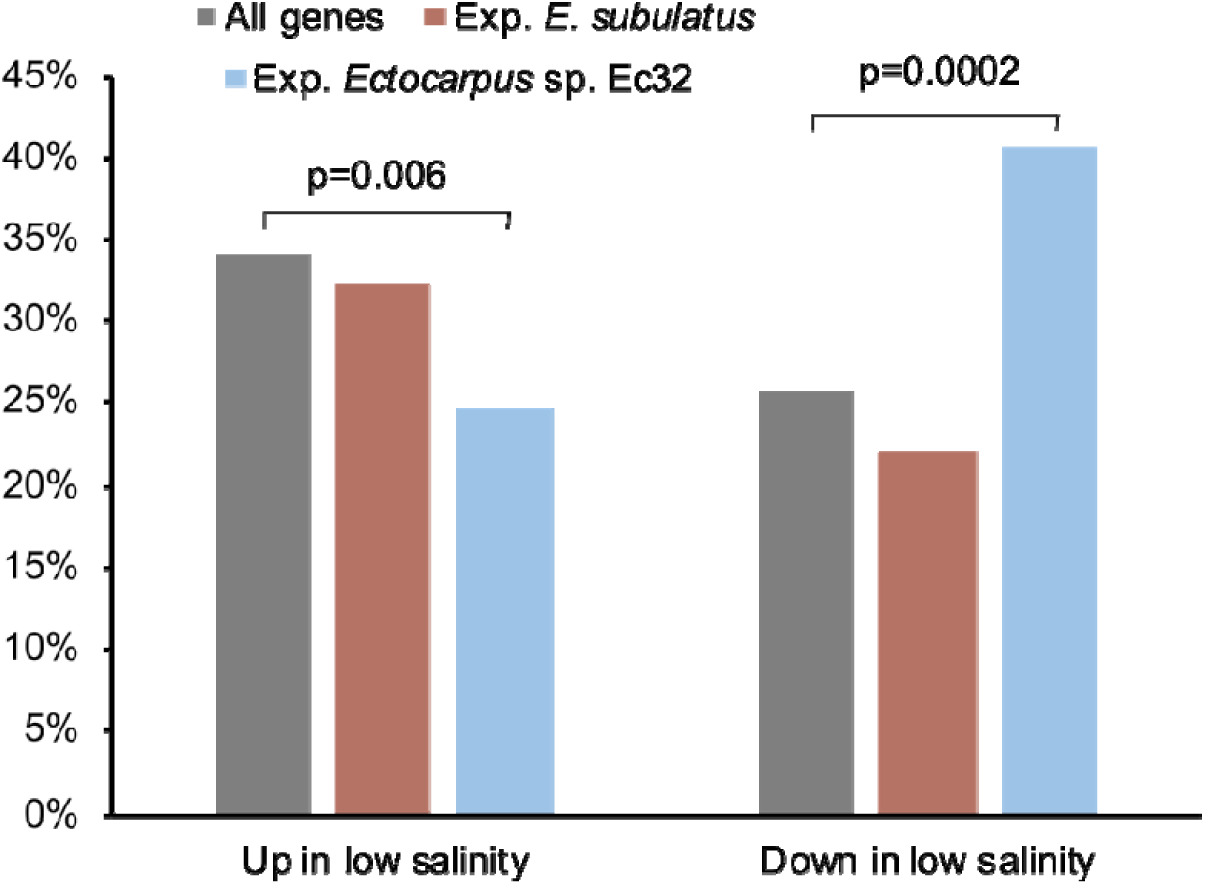
Percentage of significantly (FDR<0.05) up- and down-regulated genes in *E. subulatus* in response to low salinity (5% seawater). Grey bars are values obtained for all genes with expression data (n=6,492), while brown and blue bars include only genes belonging to gene families expanded in *E. subulatus* Bft15b (n=99) or *Ectocarpus* sp. Ec32 (n=202), respectively (“Exp.” stands for expanded). P-values correspond to the result of a Fisher exact test. Gene expression data were obtained from previous microarray experiments^29^. Please refer to Supporting information Table S6 for additional data.

### Targeted manual annotation of specific pathways

In addition to the global analyses carried out above, genes related to cell wall metabolism, sterol metabolism, polyamine and central carbon metabolism, algal defence metabolites, transporters, and abiotic stress response were manually examined and annotated, because, based on literature studies, these functions could be expected to explain the physiological differences between *E. subulatus* Bft15b and *Ectocarpus* sp. Ec32. Overall the differences between both *Ectocarpus* strains with respect to these genes were minor; a detailed description of these results is available in Supporting Information Text S1 and Supporting Information Table S7, and a brief overview of the main differences is presented below.

Regarding gene families reduced in *E. subulatus* Bft15b or expanded in *Ectocarpus* sp. Ec32, the *E. subulatus* genome encoded only 320 WSC-domain containing proteins, vs. 444 in *Ectocarpus* sp.. Many of these genes were down-regulated in response to low salinity, (61% of the WSC domain containing genes with available expression data; Fischer exact test, p=0.0004) while only 7% were upregulated (Fischer exact test, p-value=0.0036). In yeast, WSC domain proteins may act as cell surface mechanosensors and activate the intracellular cell wall integrity signalling cascade in response to hypo-osmotic shock^30^. Whether or not they have similar functions in brown algae, however, remains to be established. Furthermore, we found fewer aryl sulfotransferase, tyrosinases, potential bromoperoxidases, and thyroid peroxidases in the *E. subulatus* genome compared to *Ectocarpus* sp., and it entirely lacks haloalkane dehalogenases (Supporting Information Text S1). All of these enzymes are involved in the production of polyphenols and halogenated defence compounds, suggesting that *E. subulatus* may be investing less energy in defence, although a potential bias induced by differences in the assembly completeness cannot be excluded here.

Regarding gene families expanded in *E. subulatus* Bft15b or reduced in *Ectocarpus* sp. Ec32, we detected differences with respect to a few “classical” stress response genes. Notably an HSP20 protein was present in three copies in the genome of *E. subulatus* and only one copy in *Ectocarpus* sp.. We also found a small group of LHCX-family chlorophyll-binding proteins (CBPs) as well as a larger group belonging to the LHCF/LHCR family that have probably undergone a recent expansion in *E. subulatus* (Figure 4). Some of the proteins appeared to be truncated (marked with asterisks), but all of them were associated with RNA-seq reads, suggesting that they may be functional. A number of these proteins were also flanked by LTR-like sequences. CBPs have been reported to be up-regulated in response to abiotic stress in stramenopiles^31,32^, including *Ectocarpus*^33^, probably as a way to deal with excess light energy when photosynthesis is affected.

**Figure 4:**
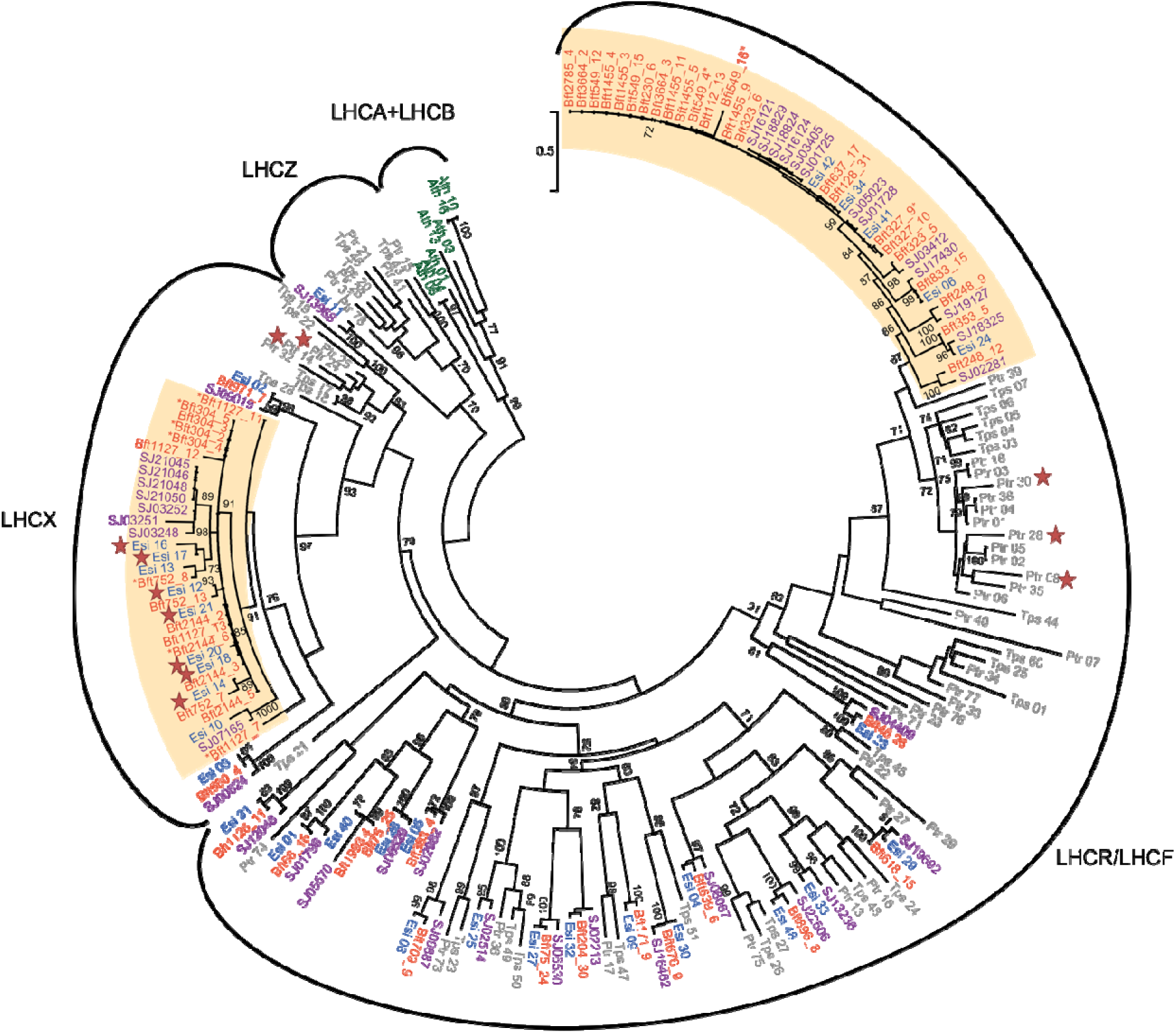
Maximum likelihood tree of chlorophyll binding proteins (CBPs) sequences in *E. subulatus* Bft15b (orange) *Ectocarpus* sp. Ec32 (blue), *S. japonica* (purple), and diatoms (*Thalassiosira pseudonana* and *Phaeodactylum tricornutum*, grey). Support values correspond to the percentage of bootstrap support from 1000 replicate runs, only values ≥ 70% are shown. *A. thaliana* sequences (green) were added as outgroup. Accessions for *E. subulatus* Bft15b are given without the Esu prefix; for *Ectocarpus* sp. Ec32, diatoms and *A. thaliana*, see^93^. Stars indicate genes that have been previously shown to be stress-induced^93^, asterisks next to the protein names indicate incomplete proteins. Probable expansions in *E. subulatus* Bft15b are indicated by an ocher background.

## Discussion

Here we present the draft genome and metabolic network of *E. subulatus* strain Bft15b, a brown alga which, compared to *Ectocarpus* sp. Ec32, is characterized by high abiotic stress tolerance^10,17^. Based on time-calibrated molecular trees, both species separated roughly 16 Mya^29^, *i.e.* slightly before the split between *Arabidopsis thaliana* and *Thellungiella salsuginea* (7-12 Mya)^34^. This split was probably followed by an adaptation of *E. subulatus* to highly fluctuating and low salinity habitats^19^.

### Traces of recent transposon activity and integration of viral sequences

The *E. subulatus* Bft15b genome is only approximately 6% (flow cytometry) to 23% (genome assembly) larger than that of *Ectocarpus* sp. Ec32, and no major genomic rearrangements or duplications were detected. However, we observed traces of recent transposon activity, especially from LTR transposons, which is in line with the absence of DNA methylation. Bursts in transposon activity have been identified as one potential driver of local adaptation and speciation in other model systems such as salmon^35^ or land plants^34,36^. Furthermore, LTRs are known to mediate the retrotransposition of individual genes, leading to the duplication of the latter^37^. In *E. subulatus* Bft15b, only a few expansions of gene families were observed since the separation from *Ectocarpus* sp. Ec32, and only in the case of the recent expansion of the LHCR family were genes flanked by a pair of LTR-like sequences. These elements lacked both the group antigen (GAG) and reverse transcriptase (POL) proteins, which implies that, if retro-transposition was the mechanism underlying the expansion of this group of proteins, it would have depended on other active transposable elements to provide these activities.

A second factor that has shaped the *Ectocarpus* genomes were viruses. Viral infections are a common phenomenon in Ectocarpales^38^, and a well-studied example is the *Ectocarpus siliculosus* virus-1 (EsV-1)^39^. It was found to be present latently in several strains of *Ectocarpus* sp. closely related to strain Ec32, and has also been found integrated in the genome of the latter, although it is not expressed^7^. As previously indicated by comparative genome hybridization experiments^40^, the *E. subulatus* Bft15b genome does not contain a complete EsV-1 like insertion, although a few shorter EsV-1-like proteins were found. Thus, the EsV-1 integration observed in *Ectocarpus* sp. Ec32 has likely occurred after the split with *E. subulatus*, and the biological consequences of this insertion remain to be explored.

### Few classical stress response genes but no transporters involved in adaptation

One aim of this study was to identify genes that may potentially be responsible for the high abiotic stress and salinity tolerance of *E. subulatus*. Similar studies on genomic adaptation to changes in salinity or to drought in terrestrial plants have previously highlighted genes generally involved in stress tolerance to be expanded in “extremophile” organisms. Examples are the expansion of catalase, glutathione reductase, and heat shock protein families in desert poplar^24^, arginine metabolism in jujube^41^, or genes related to cation transport, abscisic acid signalling, and wax production in *T. salsuginea*^34^. In our study, we found that gene families reduced in *E. subulatus* Bft15b compared to the marine *Ectocarpus* sp. Ec32 model have previously been shown to be repressed in response to stress, whereas gene families up-regulated in response to stress had a higher probability of being conserved. However, there are only few signs of known stress response gene families among them, notably the two additional HSP20 proteins and an expanded family of CBPs. *E. subulatus* Bft15b also has a slightly reduced set of genes involved in the production of halogenated defence compounds that may be related to its habitat preference: it is frequently found in brackish and even freshwater environments with low availability of halogens. It also specializes in habitats with high levels of abiotic stress compared to most other brown algae, and may thus invest less energy in defence against biotic stressors.

Another anticipated adaptation to life in varying salinities lies in modifications of the cell wall. Notably, the content of sulfated polysaccharides is expected to play a crucial role as these compounds are present in all marine plants and algae, but absent in their freshwater relatives^42,43^. The fact that we found only small differences in the number of encoded sulfatases and sulfotransferases indicates that the absence of sulfated cell-wall polysaccharides previously observed in *E. subulatus* in low salinities^44^ is probably a regulatory effect or simply related to the lack of sulfate in low salinity. This is also coherent with the wide distribution of *E. subulatus* in marine, brackish water, and freshwater environments.

Finally, transporters have previously been described as a key element in plant adaptation to different salinities^45^. Similar results have also been obtained for *Ectocarpus* in a study of quantitative trait loci (QTLs) associated with salinity and temperature tolerance^46^. In our study, however, we found no indication of genomic differences related to transporters between the two species. This observation corresponds to previous physiological experiments indicating that *Ectocarpus*, unlike many terrestrial plants, responds to strong changes in salinity as an osmoconformer rather than an osmoregulator, *i.e.* it allows the intracellular salt concentration to adjust to values close to the external medium rather than keeping the intracellular ion composition constant^33^.

### Species-specific genes of unknown function are likely to play a dominant role in adaptation

In addition to genes that may be directly involved in the adaptation to the environment, we found several gene clusters containing domains potentially involved in cell-cell signalling that were expanded in the *Ectocarpus* sp. Ec32 genome (Table 2), *e.g.* a family of ankyrin repeat-containing domain proteins^47^. These observed differences may be, in part, responsible for the existing pre-zygotic reproductive barrier between the two examined species of *Ectocarpus*^48^.

The vast majority of genomic differences between the two investigated species of *Ectocarpus,* however, corresponds to proteins of entirely unknown functions. All of the 11 gene pairs under positive selection were unknown genes taxonomically restricted to brown algae. Of the 1,611 *E. subulatus* Bft15b-specific genes, 88% were unknown. Most of these genes were expressed and are thus likely to correspond to true genes; their absence from the *Ectocarpus* sp. Ec32 genome was also confirmed at the nucleotide level. A large part of the mechanisms that underlie the adaptation to different ecological niches in *Ectocarpus* may, therefore, lie in these genes of unknown function. This can be partly explained by the fact that still only few brown algal genomes have been sequenced, and that currently most of our knowledge on the function of their proteins is based on studies in model plants, animals, yeast, or bacteria, which have evolved independently from stramenopiles for over 1 billion years^49^. They differ from land plants even in otherwise highly conserved aspects, for instance in their life cycles, cell walls, and primary metabolism^50^. Substantial contributions of lineage-specific genes to the evolution of organisms and the development of innovations have also been described for animal models^51^, and studies in basal metazoans furthermore indicate that they are essential for species-specific adaptive processes^52^.

Despite the probable importance of these unknown genes for local adaptation, *Ectocarpus* may still heavily rely on classical stress response genes for abiotic stress tolerance. Many of the gene families known to be related to stress response in land plants (including transporters and genes involved in cell wall modification), and for which no significant differences in gene contents were observed, have previously been reported to be strongly regulated in response to environmental stress in *Ectocarpus*^29,33,53^. This high transcriptomic plasticity is probably one of the features that allow *Ectocarpus* to thrive in a wide range of environments, and may form the basis for its capacity to further adapt to “extreme environments” such as freshwater^18^.

### Conclusion and future work

We have shown that since the separation of *E. subulatus* and *Ectocarpus sp.* Ec32, both genomes have been shaped partially by the activity of viruses and transposons, particularly large retrotransposons. Over this period of time, *E. subulatus* has adapted to environments with high abiotic variability including brackish water and even freshwater. We have identified a few genes that likely contribute to this adaptation, including HSPs, CBPs, a reduction of genes involved in halogenated defence compounds, or some changes in cell wall polysaccharide-modifying enzymes. However, the majority of genes that differ between the two examined *Ectocarpus* species or that may be under positive selection encode proteins of unknown function. This underlines the fundamental differences that exist between brown algae and terrestrial plants or other lineages of algae. Studies as the present one, *i.e.* without strong *a priori* assumptions about the mechanisms involved in adaptation, are therefore essential to start elucidating the specificities of this lineage as well as the various functions of the unknown genes.

## Methods

**Biological material.** Haploid male parthenosporophytes of *E. subulatus* strain Bft15b (Culture Collection of Algae and Protozoa CCAP accession 1310/34), isolated in 1978 by Dieter G. Müller in Beaufort, North Carolina, USA, were grown in 14 cm (100 ml) Petri Dishes in Provasoli-enriched seawater^54^ under a 14/10 daylight cycle at 14°C. Strains were exanimated by light microscopy (800X magnification, phase contrast) to ensure that they were free of contaminating eukaryotes, but did still contain some alga-associated bacteria. Approximately 1 g fresh weight of algal culture was dried on a paper towel and immediately frozen in liquid nitrogen. For RNA-seq experiments, in addition to Bft15b, a second strain of *E. subulatus*, the diploid freshwater strain CCAP 1310/196 isolated from Hopkins River Falls, Australia^18^, was included. One culture was grown as described above for Bft15b, and for a second culture, seawater was diluted 20-fold with distilled water prior to the addition of Provasoli nutrients^29^ (culture condition referred to as freshwater).

**Flow cytometry** experiments to measure nuclear DNA contents were carried out as previously described^55^, except that young sporophyte tissue was used instead of gametes. Samples of the genome-sequenced *Ectocarpus* sp. strain Ec32 (CCAP accession 1310/4 from San Juan de Marcona, Peru) were run in parallel as a size reference.

**DNA and RNA** were extracted using a phenol-chloroform-based protocol^56^. For DNA **sequencing**, four Illumina libraries were prepared and sequenced on a HiSeq2000: one paired-end library (Illumina TruSeq DNA PCR-free LT Sample Prep kit #15036187, sequenced with 2×100 bp read length), and three mate-pair libraries with span sizes of 3kb, 5kb, and 10kb respectively (Nextera Mate Pair Sample Preparation Kit; sequenced with 2×50bp read length). One poly-A enriched RNA-seq library was generated for each of the three aforementioned cultures according to the Illumina TruSeq Stranded mRNA Sample Prep kit #15031047 protocol and sequenced with 2×50 bp read length.

The degree **of DNA methylation** was examined by HPLC on CsCl-gradient purified DNA^56^ from three independent cultures per strain as previously described^57^.

Redundancy of mate-pairs (MPs) was reduced to mitigate the negative effect of redundant chimeric MPs during scaffolding. To this means, mate-pair reads were aligned with bwa-0.6.1 to a preliminary *E. subulatus* Bft15b draft assembly calculated from paired-end data only. Mate-pairs that did not map with both reads were removed, and for the remaining pairs, read-starts were obtained by parsing the cigar string using Samtools and a custom Pearl script. Mate-pairs with redundant mapping coordinates were removed for the final **assembly**, which was carried out using SOAPDenovo2^58^. Scaffolding was then carried out using SSPACE basic 2.0^59^ (trim length up to 5 bases, minimum 3 links to scaffold contigs, minimum 15 reads to call a base during an extension) followed by a run of GapCloser (part of the SOAPDenovo package, default settings). A dot plot of syntenic regions between *E. subulatus* Bft15b and *Ectocarpus* sp. Ec32 was generated using D-Genies 1.2.0^60^. Given the high degree of synteny observed (Supporting Information Text S1), additional scaffolding was carried out using MeDuSa and the *Ectocarpus* sp. Ec32 genome as reference^61^. This super-scaffolding method assumes that both genome structures are be similar. Annotations were generated first for version 1 of the Bft15b genome and then transferred to the new scaffolds of version 2 using the ALLMAPS^62^ liftover function. Both the assemblies with (V2) and without (V1) MeDuSa scaffolding have been made available. RNA-seq reads were cleaned using Trimmomatic (default settings), and a second Bft15b genome-guided assembly was performed with Tophat2 and with Cufflinks. Sequencing coverage was calculated based on mapped algal reads only, and for mate-pair libraries the genomic coverage was calculated as number of unique algal mate-pairs * span size / assembly size.

As cultures were not treated with antibiotics prior to DNA extraction, **bacterial scaffolds were removed** from the final assembly using the taxoblast pipeline^63^. Every scaffold was cut into fragments of 500 bp, and these fragments were aligned (blastn, e-value cutoff 0.01) against the GenBank non-redundant nucleotide (nt) database. Scaffolds for which more than 90% of the alignments were with bacterial sequences were removed from the assembly (varying this threshold between 30 and 95% resulted in only very minor differences in the final assembly). Finally, we ran the Anvi’o v5 pipeline to identify any remaining contaminant bins (both bacterial and eukaryote) based on G/C and kmer contents as well as coverage^64^. “Contaminant” scaffolds were submitted to the MG-Rast server to obtain an overview of the taxa present in the sample^65^. They are available at http://application.sb-roscoff.fr/blast/subulatus/download.html.

**Repeated elements** were searched for *de novo* using TEdenovo and annotated using TEannot with default parameters. LTR-like sequences were predicted by the LTR-harvest pipeline^66^. These tools are part of the REPET pipeline^67^, of which version 2.5 was used for our dataset.

**BUSCO** 2.0 analyses^27^ were run on the servers of the IPlant Collaborative^68^ with the general eukaryote database as a reference and default parameters and the predicted proteins as input.

**Plastid and mitochondrial genomes** of *E. subulatus* Bft15b, were manually assembled based on scaffolds 416 and 858 respectively, using the published organellar genomes of *Ectocarpus* sp. Ec32 (accessions NC_013498.1, NC_030223.1) as a guide^7,69,70^. Genes were manually annotated based on the result of homology searches with *Ectocarpus* sp. Ec32 using a bacterial genetic code (11) and based on ORF predictions using ORF finder. Ribosomal RNA sequences were identified by RNAmmer^71^ for the plastid and MITOS^72^ for the plastid, and tRNAs or other small RNAs were identified using ARAGORN^73^ and tRNAscan-SE^74^. In the case of the mitochondrial genome, the correctness of the manual assembly was verified by PCR where manual and automatic assemblies diverged.

Putative **protein-coding sequences** were identified using Eugene 4.1c^75^. Assembled RNA-seq reads were mapped against the assembled genome using GenomeThreader 1.6.5, and all available proteins from the Swiss-Prot database as well as predicted proteins from the *Ectocarpus* sp. Ec32 genome^7^ were aligned to the genome using KLAST^76^. Both aligned *de novo*-assembled transcripts and proteins were provided to Eugene for gene prediction, which was run with the parameter set previously optimized for the *Ectocarpus* sp. Ec32 genome^7^. The subcellular localization of the proteins was predicted using SignalP version 4.1^77^ and the ASAFIND software version 1.1.5^26^.

**For functional annotation**, predicted proteins were submitted to InterProScan and compared to the Swiss-Prot database by BlastP search (e-value cutoff 1e-5), and the results imported to Blast2GO^78^ The genome and all automatic annotations were imported into Apollo^79,80^ for manual curation. During manual curation sequences were aligned with characterized reference sequences from suitable databases (e.g. CAZYME, TCDB, SwissProt) using BLAST, and the presence of InterProScan domains necessary for the predicted enzymatic function was manually verified.

The *E. subulatus* Bft15b **genome-scale metabolic model** reconstruction was carried out as previously described^28^ by merging an annotation-based reconstruction obtained with Pathway Tools^81^ and an orthology-based reconstruction based on the *Arabidopsis thaliana* metabolic network AraGEM^82^ using Pantograph^83^. A final step of gap-filling was then carried out using the Meneco tool^84^. The entire reconstruction pipeline is available via the AuReMe workspace^85^. For pathway-based analyses, pathways that contained only a single reaction or that were less than 50% complete were not considered.

**Functional comparisons of gene contents** were based primarily on orthologous clusters of genes shared with version 2 of the *Ectocarpus* sp. Ec32 genome^86^ as well as the *S. japonica* (Areschoug) genome^11^. They were determined by the OrthoFinder software version 0.7.1^87^. To identify genes specific to either of the *Ectocarpus* genomes, we examined all proteins that were not part of a multi-species cluster and verified their absence in the other genome by tblastn searches (threshold e-value of 1e-10). Only genes without tblastn hit that encoded proteins of at least 50 amino acids were further examined. A second approach consisted in identifying clusters of genes that were expanded or reduced in either of the two *Ectocarpus* genomes based on the Orthofinder results. Blast2GO 3.1^78^ was then used to identify significantly enriched GO terms among the genes specific to either *Ectocarpus* genome or the expanded/reduced gene families (Fischer’s exact test with FDR correction FDR<0.05). These different sets of genes were also examined manually for function, genetic context, GC content, and EST coverage (to ensure the absence of contaminants).

The search for **genes under positive selection** was based on a previous analysis in other brown algae^88^. Therefore, Orthofinder analyses were expanded to include also *Macrocystis pyrifera, Scytosiphon lomentaria*^88^, and *Cladosiphon okamuranus*^12^. Rates of non-synonymous to synonymous substitution (ω=dN/dS) were searched for in clusters of single-copy orthologs. Protein sequences were aligned with Tcoffee^89^ (M-Coffee mode), translated back to nucleotide using Pal2Nal^90^, and curated with Gblocks^91^ (-t c -b4 20) or manually when necessary. Sequences that produced a gapless alignment that exceeded 100bp were retained for pairwise dN/dS analysis between *Ectocarpus* strains using CodeML (F3×4 model of codon frequencies, runmode = −2) of the PAML4 suite^92^. Orthogroups for which the pairwise dN/dS ratio between *Ectocarpus* species exceeded 0.5, which were not saturated (dS < 1), and which contained single-copy orthologs in at least two other species were used to perform positive selection analysis with CodeML (PAML4, F3×4 model of codon frequencies): branch-site models were used to estimate dN/dS values by site and among branches in the species tree generated for each orthogroup. The branch leading to the genus *Ectocarpus* was selected as a ‘foreground branch’, allowing different values of dN/dS among sites in contrast to the remaining branches that shared the same distribution of ω. Two alternative models were tested for the foreground branch: H1 allowing the dN/dS to exceed 1 for a proportion of sites (positive selection), and H0 constraining dN/dS<1 for all sites (neutral and purifying selection). A likelihood ratio test was then performed for the two models (LRT=2×(lnLH1-lnLH0)) and genes for which H1 fitted the data significantly better (p<0.05) were identified as evolving under positive selection.

**Phylogenetic analyses** were carried out for gene families of particular interest. For chlorophyll-binding proteins (CBPs), reference sequences were obtained from a previous study^93^, and aligned together with *E. subulatus* Bft15b and *S. japonica* CBPs using MAFFT (G-INS-i)^94^. Alignments were then manually curated, conserved positions selected in Jalview^95^, and maximum likelihood analyses carried out using PhyML 3.0^96^, the LG substitution model, 1000 bootstrap replicates, and an estimation of the gamma distribution parameter. The resulting phylogenetic tree was visualized using MEGA7^97^.

## Acknowledgements

We would like to thank Philippe Potin, Mark Cock, Susanna Coelho, Florian Maumus, and Olivier Panaud for helpful discussions, as well as Gwendoline Andres for help setting up the Jbrowse instance. This work was funded partially by ANR project IDEALG (ANR-10-BTBR-04) “Investissements d’Avenir, Biotechnologies-Bioressources”, the European Union’s Horizon 2020 research and innovation Programme under the Marie Sklodowska-Curie grant agreement number 624575 (ALFF), and the CNRS Momentum call. Sequencing was performed at the Genomics Unit of the Centre for Genomic Regulation (CRG), Barcelona, Spain.

## Author contributions

Conceived the study: SMD, AP, AS, HH, CB, TT. Provided materials: AFP, APL. Performed experiments: SMD, SD, IGN, DM, MMP. Analysed data: SMD, APL, EC, LBG, NP, MA, KA, CHC, JC, AC, LD, SD, CF, AGo, AGr, CH, DJ, HK, XL, GVM, AEM, MM, PP, MMP, ASim, CT, HSY, TT. Wrote the manuscript: SMD, KA, APL, JC, LD, CH, Ago, AGr, GVM, ASim, TT. Revised and approved of the final manuscript: all authors.

## Additional Information

### Data availablility

Sequence data (genomic and transcriptomic reads) were submitted to the European Nucleotide Archive (ENA) under project accession number PRJEB25230 using the EMBLmyGFF3 script^98^. A JBrowse^99^ instance comprising the most recent annotations is available via the server of the Station Biologique de Roscoff (http://mmo.sb-roscoff.fr/jbrowseEsu). The reconstructed metabolic network of *E. subulatus* is available at http://gem-aureme.irisa.fr/sububftgem. Additional resources and annotations including a blast server are available at http://application.sb-roscoff.fr/project/subulatus/index.html. The complete set of manual annotations is provided in Supporting Information Table S7.

