## Supplementary Text for "The genome of *Ectocarpus subulatus* – a highly stress-tolerant brown alga"

### Synteny analyses

D-Genies^1^ analyses revealed an overall high degree of synteny between the two *Ectocarpus* genomes (diagonal line in Figure S1), with some minor variations, but without major gene shuffling or duplications.

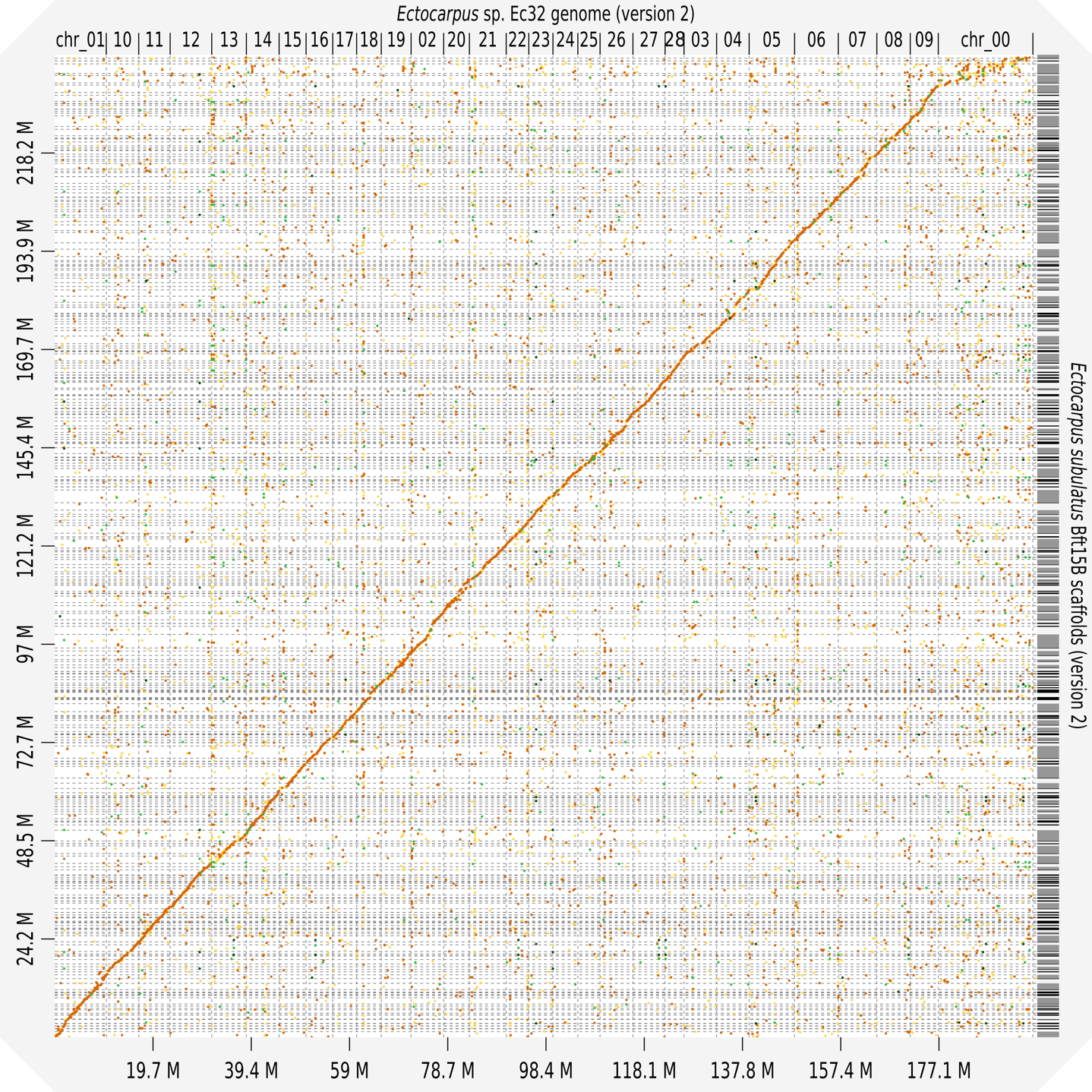

**Figure S1**: Synteny plot for the comparison of *E. subulatus* Bft15b scaffolds with *Ectocarpus* sp. Ec32 generated using D-Genies. Colors represent the percentage of identity between anchors (yellow=0-25%; orange=25-50%, light green=50-75; dark green=75-100%. An identical figure was obtained for Version 1 of the *E. subulatus* genome.

### Organellar genomes

The mitochondrial genome of *E. subulatus* Bft15b differed from that of *Ectocarpus* sp. Ec32 essentially with respect to the presence of four additional introns, three located within the 16S and 23S rRNA genes and including a hypothetical maturase (Figure S2). It is the presence of these introns that is responsible for the ca. 8 kb size difference compared to the mitochondrial genome of *Ectocarpus* sp. Ec32.

A large structural difference was observed only in the plastid genome where one inversion of *ca.* 50 kb in the small single copy (SSC) region may have occurred. Furthermore, small differences in gene contents of the *E. subulatus* Bft15b plastid with respect to *Ectocarpus* sp. Ec32 were detected around two inverted repeat (IR) regions concerning the following genes: *psbC* (gene truncated), *psbD* (IR region next to gene), *rpoB* (large gap, frameshift), and *tRNA-Arg* and *tRNA-Glu* (duplicated in the tRNA region). Pseudogenization of genes at the edge of IRs is indeed a common phenomenon^2^.

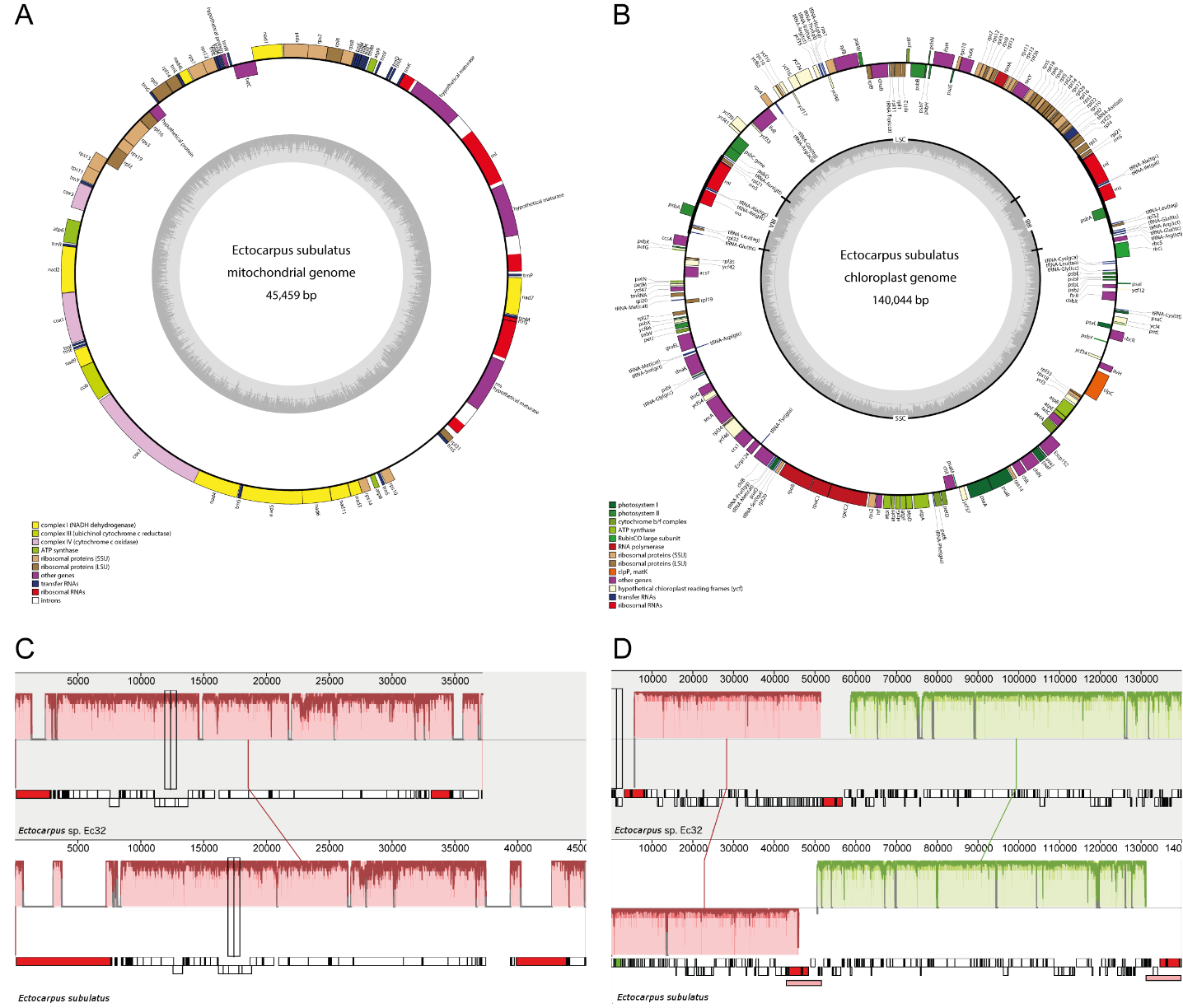

**Figure S2**: Mitochondrial and plastidial genomes of *E. subulatus* Bft15b visualized using OrganellarGenomeDRAW^3^ (panels A and B), and aligned with the *Ectocarpus* sp. Ec32 organelles using Mauve 2.3.1^4^ (panels C and D). In panels C and D blocks of the same color correspond to orthologous sequences.

### Manually annotated gene families

#### Cell wall metabolism

Cell walls are key components of both plants and algae and, as a first barrier to the surrounding environment, important for many processes including development and the acclimation to environmental changes. Synthesis and degradation of cell wall oligo- and polysaccharides is facilitated by carbohydrate-active enzymes (CAZymes)^5^. These comprise several families including glycoside hydrolases (GHs) and polysaccharide lyases (PLs), both involved in the cleavage of glycosidic linkages, glycosyltransferases (GTs), which create glycosidic linkages, and additional enzymes such as the carbohydrate esterases (CEs) which remove methyl or acetyl groups from substituted polysaccharides.

The genome of *E. subulatus* Bft15b encodes 37 GHs (belonging to 17 GH families), 94 GTs (belonging to 28 GT families), nine sulfatases (family S1-2), and 13 sulfotransferases, but lacks genes homologous to known PLs and CEs (Table S8). In terms of the number of CAZY families, similar results were obtained for E*. subulatus* Bft15b compared to the *Ectocarpus* sp. Ec32 and *S. japonica* genomes (Cock *et al.*, 2010; Ye *et al.*, 2015). In terms of the total CAZY gene number, the results are slightly lower in *E. subulatus* Bft15b compared to the other two genomes. Interestingly, *S. japonica* features an expansion of certain CAZY families probably related to the establishment of more complex tissues in this kelp (*i.e.* 82 GHs belonging to 17 GH families in *E. subulatus* Bft15b, and 131 GTs belonging to 31 GT families in *S. japonica*).

As *Ectocarpus* sp. Ec32 and *S. japonica*, *E. subulatus* Bft15b possesses CAZY families that are absent from green plants but present in other phyla such as bacteria (GH88), animals (GT23 and GT49), and *Amoebozoa* (GT60). Some variations in the GT families were observed between the three genomes with some families present in only one of the brown algal genomes (*e.g.* GH30, GT15, GT18, GT25, GT28, GT50, GT54, GT65, GT66, GT74, GT77). In these cases, the number of members per family was low (*e.g.* the GT24 family has one member in *Ectocarpus* sp. Ec32, two in *E. subulatus* Bft15b and none in *S. japonica*) and the encoded proteins frequently belonged to GT families only poorly defined. These findings are therefore to be taken cautiously, and the specific metabolisms or activities in which such genes are involved are difficult to predict.

Brown algal cell walls are composed predominantly of polyanionic polysaccharides, mainly alginates and fucose-containing sulfated polysaccharides^6,7^. These polymers are prevalent over neutral and crystalline components, which include cellulose. The cell wall biosynthesis likely follows the same pathways in the three brown algae investigated and putative genes have been previously reported for *Ectocarpus* sp. Ec32^8^. Overall, the numbers of genes putatively involved in alginate, fucan, and cellulose synthesis are within the same range between the two Ectocarpales and *S. japonica*, except an expansion of GT2s and of mannuronan C5-epimerases (ManC5-Es) likely to catalyze the terminal steps of alginate synthesis^9,10^ in *S. japonica* (*i.e.* 28, 24 and 105 ManC5-E in *Ectocarpus* sp. Ec32, *E. subulatus* Bft15b, and *S. japonica*, respectively).

As in land plants, cellulose in brown algae is likely to be synthesized by genes of the GT2 family. Among the nine GT2 enzymes identified in the *E. subulatus* Bft15b genome, eight are homologous to cellulose synthases and cellulose synthase-like proteins. One cellulose synthase in *E. subulatus* Bft15b (EsuBft84_3) has been duplicated in *Ectocarpus* sp. Ec32 (Esi0097_0024, Esi0097_0016). This gene family is expanded in *S. japonica* with 24 GT2s. We also found a homolog of the first characterized brown algal alginate lyase from *S. japonica* in *E. subulatus* Bft15b (EsuBft565_4). This gene is likely involved in the active remodeling of cell walls algal growth and development. However, the three brown algal genomes investigated, known families of cellulases and expansins are absent, suggesting the existence of novel mechanisms and⁄or of novel proteins for cell wall expansion in brown algae as compared to land plants.

Genes homologous to the D-4,5 unsaturated β-glucuronyl hydrolase (GH88 family) were found, with one gene in *E. subulatus* Bft15b (EsuBft2077_2) being appended to a ManC5-E. GH88 may act specifically on unsaturated oligosaccharides released by polysaccharide lyases, indicating that brown algae may possess alginate lyases belonging to novel PL families.

Regarding fucan metabolism, several GT families (GT10, GT23, GT65) were previously discussed to be a source of candidates for the polymerization of GDP-fucose into the elongating fucan chain^8^. No GT65 genes were identified either in *S. japonica* or *E. subulatus* Bft15b (Table S8), the GT10 family is small in all the three genomes (one or two genes), and the GT23 family shows an expansion in *S. japonica* (2 genes in the two Ectocarpales and 17 genes in *S. japonica*).

The species *E. subulatus* is frequently found in brackish- and even freshwater environments^11^ where its cell wall exhibits little or no sulfation^12^. Hence, we also assessed whether *E. subulatus* Bft15b had reduced the gene families responsible for this process*.* Its genome encodes only eight sulfatases and six sulfotransferases compared to ten and seven, respectively, in *Ectocarpus* sp. Ec32.

#### Sterol metabolism

Sterols are modulators of membrane fluidity among eukaryotes, and provide the backbone for signalling molecules^13^. The sterol content and composition of cell membranes has previously been related to differences in salinity tolerance, *e.g.* in some aquatic yeasts^14^. Fucosterol, cholesterol, and ergosterol are the most abundant sterols in *Ectocarpus* sp. Ec32, where their relative abundance varies according to sex and temperature^15^. All three molecules are thought to be synthesized from squalene by a succession of 12 to 14 steps, relying on a roughly conserved set of twelve enzymes^13^. The *E. subulatus* Bft15b and *Ectocarpus* sp. Ec32 genomes each encode homologs of twelve of them (squalene monooxygenase, oxydosqualene cyclase, sterol C14 demethylase, sterol C14 reductase, sterol C4 methyloxidase, hydroxysteroid 3β-dehydrogenase, sterol-delta(8), delta7-isomerase, cyclopropylsterol isomerase, sterol delta-7 reductase, sterol C5 desaturase, and two sterol methyltransferases). The remaining two, a delta-24-reductase (DHCR24) and a C22 desaturase (CYP710), were probably lost secondarily. In land plants, these latter enzymes are involved in the two steps transforming fucosterol into stigmasterol. Fucosterol is the main sterol in brown algae, and provides a substrate for saringosterol, a brown alga-specific C24-hydroxylated fucosterol-derivative with antibacterial activity^16^.

#### Algal defences: metabolism of phenolics and halogens

It is generally assumed that organisms need to strike a balance between abiotic stress tolerance, biotic stress tolerance, and growth^17^. In particular in the intertidal zone, the upper levels are generally governed by organisms with very high abiotic stress tolerance, whereas in lower zones these organisms are outcompeted by species with higher biotic stress tolerance^18^. Considering that *E. subulatus* is a highly abiotic stress-tolerant species compared to other *Ectocarpus* species, we examined if this capacity was correlated with a reduction of genes involved in algal defences, focusing on polyphenols and halogenated compounds.

Polyphenols are a group of defence compounds in brown algae that are likely to be important both for abiotic^19^ and biotic stress tolerance^20^. Brown algae produce specific polyphenols called phlorotannins, which are analogous to land plant tannins. These products are polymers of phloroglucinol, which are synthesized via the activity of a phloroglucinol synthase, a type III polyketide synthase characterized in *Ectocarpus* sp. Ec32^21^. In analogy to the flavonoid pathway of land plants, the further metabolism of phlorotannins is thought to be driven by members of chalcone isomerase-like (CHIL), aryl sulfotransferase (AST), flavonoid glucosyltransferase (FGT), flavonoid O-methyltransferase (OMT), polyphenol oxidase (POX), and tyrosinase (TYR) families^22^. While copy numbers between the two *Ectocarpus* species and *S. japonica* are identical for PKS III, CHIL, FGT, OMT and POX, *E. subulatus* Bft15b encodes fewer ASTs and TYRs (Table S8). In the case of ASTs, this may be related to the lower concentration of sulphate in low salinity environments frequently colonized by *E. subulatus*.

Halogenated defence compounds are produced in brown algae via the activity of halogenating enzymes, *e.g.* the vanadium-dependent haloperoxidase (vHPO). While *S. japonica* has recently been reported to possess 17 potential bromoperoxidases (vBPO) and 59 putative iodoperoxidases (vIPO)^23^, *Ectocarpus* sp. Ec32 and *E. subulatus* Bft15b possess only a single vBPO each and no vIPO, but have in turn slightly expanded a haloperoxidase family closer to vHPOs characterized in several marine bacteria^24^ (Table S8). One difference between the two *Ectocarpus* species is that *E. subulatus* Bft15b possesses only three vHPO genes compared to the five copies found in the genome of Ec32. In addition, homologs of thyroid peroxidases (TPOs) may also be involved in halide transfer and biotic stress response. Again, Ec32 and Bft15b show a reduced set of these genes compared to *S. japonica*, and Ec32 contains more copies than Bft15b. Finally, a single haloalkane dehalogenase (HLD) was found exclusively in *Ectocarpus* sp. Ec32.

**Table S8:** Prevalence of several genes families related to key metabolic pathways in brown algae. *E.* sp., *Ectocarpus sp.* Ec32; *E. su*, *Ectocarpus subulatus* Bft15b*;* *S. ja*, *Saccharina japonica*; GT, glycosyltransferase; GH, glycosyl hydrolase; vBPO, vanadium-dependent bromoperoxidase; vIPO, vanadium-dependent ioperoxidase; vHPO, vanadium-dependent haloperoxidase; TPO, homolog of thyroid peroxidase; HLD, haloalkane dehalogenase; MPI, mannose-6-phosphate isomerase; PMM, phosphomannomutase; MPG, mannose-1-phosphate guanyltransferase; GMD, GDP-mannose 6-dehydrogenase; MS, mannuronan synthase; ManC5-E, mannuronan C5-epimerase; FK, L-fucokinase; GFPP, GDP-fucose pyrophosphorylase; GM46D, GDP-mannose 4,6-dehydratase; GFS, GDP-L-fucose synthetase; ST, sulfotransferase; PKS III, type III polyketide synthase; CHIL, chalcone isomerase-like; AST, aryl sulfotransferase; FGT, flavonoid glucosyltransferase; OMT, flavonoid O-methyltransferase (OMT); POX, polyphenol oxidase; TYR, tyrosinase. *for GT31, three genes have been identified in *Ectocarpus* sp. Ec32 in the present study that have not been initially annotated in the corresponding genome ^22^ and companion paper ^8^. n. i.: The presence of sulfatases in the *S. japonica* genome was not indicated in the corresponding paper ^23^.

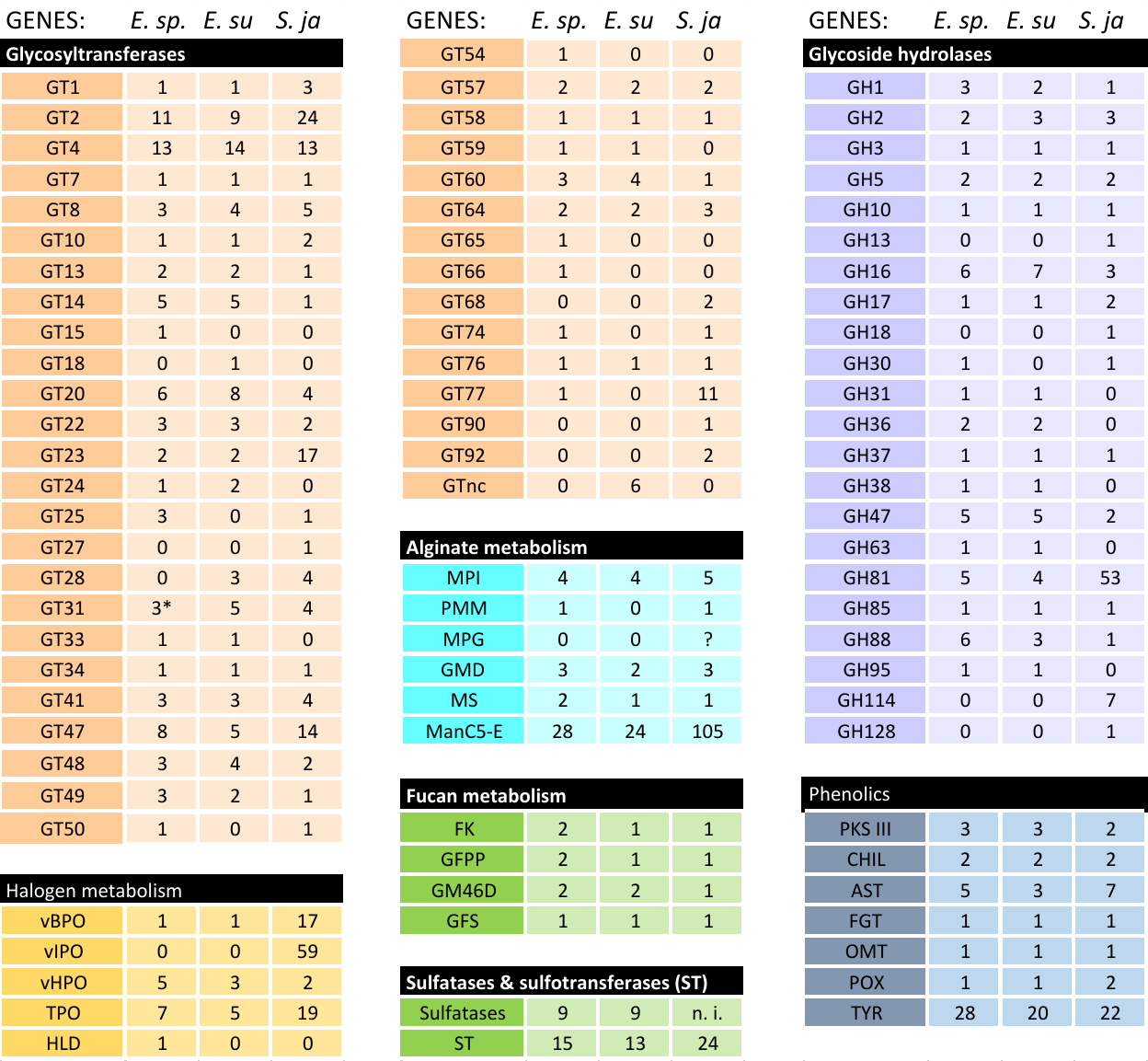

#### Transporters

Transporters are key actors driving salinity tolerance in terrestrial plants^25^. We therefore carefully assessed potential differences in this group of proteins that may explain physiological differences between Ec32 and Bft15b. Analyses were based on the five main categories of transporters described in the Transporter Classification Database (TCDB)^26^: channels/pores, electrochemical potential-driven transporters, primary active transporters, group translocators, and transmembrane electron carriers. A total of 292 genes were identified in *E. subulatus* Bft15b (Supporting Information Table S7). They consist mainly of transporters belonging to the three first categories listed above. All 27 annotated transporters of the channels/pores category belong to the alpha-type channel (1.A.) and are likely to be involved in movements of solutes by energy-independent processes. One hundred and forty-five proteins were found to correspond to the second category (electrochemical potential-driven transporters) containing transporters using a carrier-mediated process to catalyse uniport, antiport, or symport. The most represented superfamilies are APC (Amino Acid-Polyamine-Organocation, 24), DMT (Drug/Metabolite Transporter, 16), MFS (Major Facilitator Superfamily, 32), and MC (Mitochondrial Carrier, 34). Primary active transporters (third category) use a primary source of energy to drive the active transport of a solute against a concentration gradient. Eighty proteins representing this category were found in the *E. subulatus* Bft15b genome, including 59 ABC transporters and 15 belonging to the P-type ATPase superfamily. No homologs of group translocators or transmembrane electron carriers were identified, but 14 transporters were classified as category 9, which is poorly characterized. A 1:1 ratio of orthologous genes coding for all of the transporters described above was observed between both *Ectocarpus* genomes, except for EsuBft583_3, an anion-transporting ATPase, which is also present in diatoms and *S. japonica*, but may have been recently lost in *Ectocarpus* sp. Ec32.

#### Abiotic stress-related genes

Reactive oxygen species (ROS) scavenging enzymes, including ascorbate peroxidases, superoxide dismutases, catalases, catalase peroxidases, glutathione reductases, (mono)dehydroascorbate reductases, and glutathione peroxidases are important for the redox equilibrium of organisms^27^. An increased reactive oxygen scavenging capacity has been correlated with stress tolerance in brown algae^28^. In the same vein, chaperone proteins including heat shock proteins (HSPs), calnexin, calreticulin, T-complex proteins, and tubulin-folding co-factors are important for protein re-folding under stress. The transcription of these genes is very dynamic and generally increases in response to stress in brown algae^29,30^. In total, 104 genes encoding members of the protein families listed above were manually annotated in the *E. subulatus* Bft15b genome (Supporting Information Table S7) and HSP20 proteins were present in three copies in Bft15b vs. one copy in Ec32 (Table S8). Additional information regarding the number of genes related to reactive oxygen species scavenging, heat shock proteins, T-complex proteins, and tubulin folding protein found in the *E. subulatus* Bft15b and *Ectocarpus* sp. Ec32 genomes is provided in Table S9 below. Corresponding gene accession numbers can be found in Supporting Information Table S1.

**Table S9:** Comparison of known stress-response genes in *E. subulatus* Bft15b (*E. su.)* and *Ectocarpus* sp. Ec32 (*E.* sp.). For more details on the *E. subulatus* genes please refer to Supporting information Table S7; for more information on the *Ectocarpus* sp. Ec32 sequences please refer to Cock et al.^22^ or the corresponding genome database: <https://bioinformatics.psb.ugent.be/orcae/overview/EctsiV2>

| Gene | *E. su.* | *E.* sp. |
| --- | --- | --- |
| Reactive oxygen scavenging enzymes |  |  |
| ascorbate peroxidase (APX) | 2 | 3 |
| Cu/Zn superoxide dismutase (SOD) | 1 | 2 |
| Fe superoxide dismutase (SOD) | 4 | 6 |
| catalase (CAT) | 1 | 1 |
| Catalase-peroxidase | 8 | 7 |
| glutathione reductase (GR) | 2 | 3 |
| monodehydroascorbate reductase (MDHAR) | 1 | 2 |
| dehydroascorbate reductase (DHAR) | 1 | 1 |
| glutathione peroxidase (GPX) | 3 | 6 |
| heat shock proteins |  |  |
| HSP100/ClpB | 1 | 2 |
| HSP90/HtpG | 4 | 4 |
| HSP70/DnaK family^1^ | ~21 | ~21 |
| HSP60/GroEL | 2 | 3 |
| HSP40/DnaJ family^1^ | ~34 | ~41 |
| HSP20/small HSP | 3 | 1 |
| HSP10/GroES | 0 | 1 |
| Calnexin | 1 | 1 |
| Calreticulin | 1 | 1 |
| T-complex proteins |  |  |
| alpha | 1 | 1 |
| beta | 1 | 1 |
| gamma | 1 | 1 |
| delta | 1 | 1 |
| epsilon | 1 | 1 |
| zeta | 1 | 1 |
| eta | 1 | 1 |
| theta | 1 | 1 |
| tubulin folding cofactors |  |  |
| A | 1 | 1 |
| B | 1 | 1 |
| C | 1 | 1 |
| D | 1 | 1 |
| E | 1 | 1 |

#### Polyamines

Polyamines are aliphatic amines with two or more primary amine groups which are able to interact with a wide range of macromolecules, including proteins, DNA, RNA, and phospholipids. They are essential for cellular growth, regulating cell division, and maintaining normal physiological function. There are also involved in responses to multiple abiotic stresses^31–38^, and have been shown to play an important role as signal molecules controlling the transition from vegetative to reproductive development in seaweeds ^39,40^. The in-depth annotation of their biosynthetic pathway is still lacking in macroalgae. Here we have performed these annotations and confirm that both *Ectocarpus* sp. Ec32 and *E. subulatus* Bft15b possess all the genes involved in the polyamine pathway. However, no distinctive features relative to the specificities of the two species have been observed.

#### Central carbon metabolism

A characteristic feature of brown algae is that they store carbohydrates not as glycogen or starch, like most animals and plants, but as laminarin^41^. Brown algae also have the particularity of using the photoassimilate D-fructose 6-phosphate to produce the alcohol sugar D-mannitol instead of sucrose like land plants. The *E. subulatus* Bft15b genome contains similar sets of genes for carbon storage compared to *Ectocarpus* sp. Ec32: all the genes encoding enzymes involved in sucrose metabolism and starch biosynthesis are completely absent while all genes necessary for trehalose synthesis, as well as laminarin synthesis and recycling were found. Based on analysis of transcriptomic data obtained for a range of brown algae through the 1000 plants project (OneKP; <https://sites.google.com/a/ualberta.ca/onekp/>), it has also been suggested that Ectocarpales may contain three copies of M1PDH genes^42^. In contrast, only two genes were identified in transcriptomic resources of Dictyotales, Desmarestiales, Laminariales, and Fucales. Analysis of genomic resources supported this hypothesis: both *Ectocarpus* species, as well as *C. okamuranus* and *N. decipiens*, contain two genes probably resulting from a specific duplication within this order of brown algae (M1PDH1 and M1PDH2, numbering based on *Ectocarpus* sp. Ec32 data presented in^43^). Conversely, the *S. japonica* genome contains orthologs only for M1PDH1 and M1PDH3 (Figure S3). Further analysis of Phaeophyceae genomes will be appropriate to test such hypothesis.

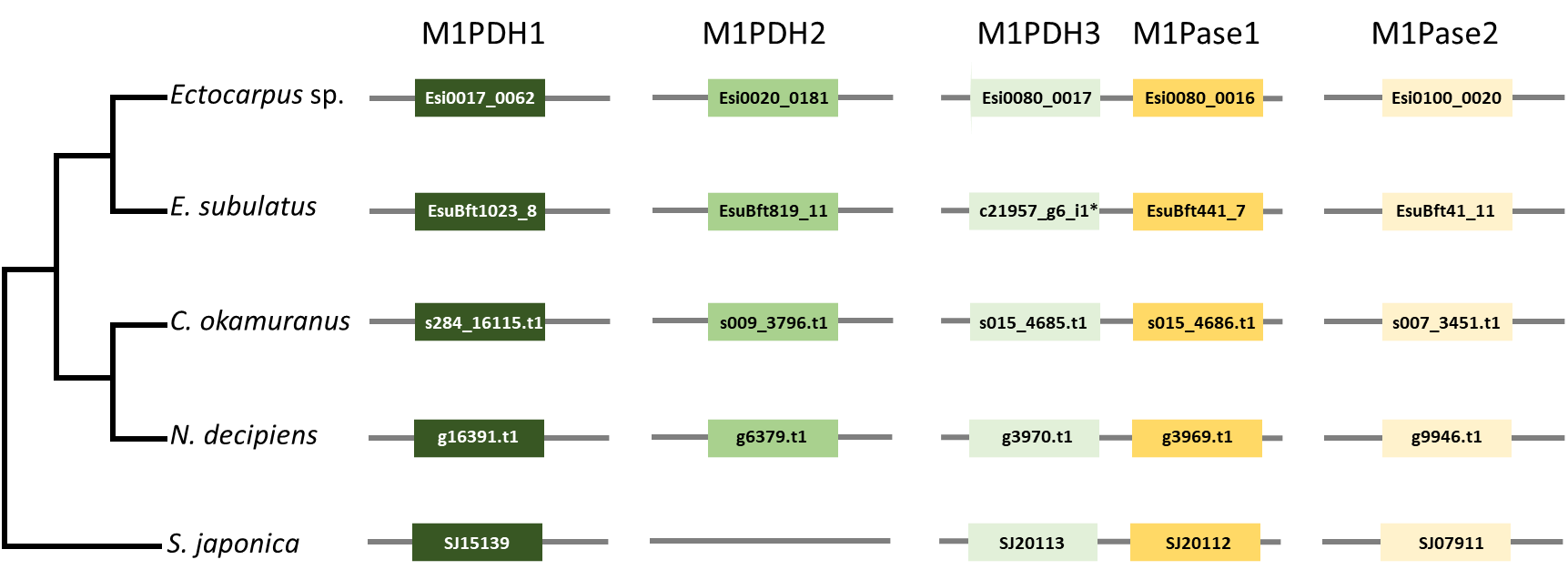

**Figure S3.** Schematic representation of candidate genes involved in mannitol synthesis in genomes of brown algae. M1PDH orthologs and paralogs are indicated in green, M1Pase orthologs and paralogs in yellow. For the *C. okamuranus* genes, the prefix “Cok_S_” has been deleted to improve clarity. * indicates a gene absent in the genome of *E. subulatus*, but identified in transcriptomic data obtained for this alga in the course of the reported study.

### Metabolic network-based comparisons

In total, the *E. subulatus* Bft15b metabolic network comprised 2,445 genes associated with 2,074 metabolic reactions and 2,173 metabolites in 464 pathways, 259 of which were complete. These results are similar to data previously obtained for *Ectocarpus* sp. Ec32 (1,977 reactions, 2,132 metabolites, 2,281 genes, 459 pathways, 272 complete pathways)^44^. Comparisons between both networks were carried out on a pathway level (Table S10), focusing on pathways present (*i.e.* complete to more than 50%) in one of the species, but with no reactions in the other. This led to the identification of 16 pathways potentially specific to *E. subulatus* Bft15b, and 11 specific to *Ectocarpus* sp. Ec32, which were further manually investigated (Table S10). In all of the examined cases, the observed differences were due to protein annotation, but not due to the presence/absence of proteins associated with these pathways in both species. For instance, the pathway "spermine and spermidine degradation III" (PWY-6441) was only found in *E. subulatus* Bft15b because the corresponding genes had been manually annotated in this species, while this was not the case in *Ectocarpus* sp. Ec32. On the other hand, three pathways related to methanogenesis (PWY-5247, PWY-5248, and PWY-5250) were falsely included in the metabolic network of *E. subulatus* Bft15b due to an overly precise automatic GO annotation of the gene Bft140_7. All in all, based on our network comparisons, we confirmed no differences regarding the presence or absence of known metabolic pathways in the two examined species of *Ectocarpus*.

**Table S10.** Comparison of metabolic pathways found in *E. subulatus* Bft15b (*E. su.)* and *Ectocarpus* sp. Ec32 (*E.* sp.) in terms of the number of reactions found. For details regarding each of the pathways please refer to the online representation of the network at: <http://gem-aureme.irisa.fr/sububftgem>.

| **Pathway** | **Total** | ***E.* sp.** | ***E. su.*** |  | **Pathway** | **Total** | ***E.* sp.** | ***E. su.*** |
| --- | --- | --- | --- | --- | --- | --- | --- | --- |
| PWY-6441 | 2 | 0 | 2 |  | PWY-5667 | 4 | 4 | 4 |
| PWY-5250 | 2 | 0 | 2 |  | PWY-6365 | 4 | 4 | 4 |
| PWY-5248 | 2 | 0 | 2 |  | ARGSYN-PWY | 4 | 4 | 4 |
| PWY-723 | 2 | 0 | 2 |  | VALSYN-PWY | 4 | 4 | 4 |
| PWY-6754 | 2 | 0 | 2 |  | PWY0-1415 | 4 | 4 | 4 |
| PWY-5874 | 2 | 0 | 2 |  | PWY4FS-6 | 4 | 4 | 4 |
| PWY0-501 | 2 | 0 | 2 |  | PWY-7197 | 4 | 4 | 4 |
| PWY-5247 | 2 | 0 | 2 |  | PWY-7279 | 4 | 4 | 4 |
| PWY-5311 | 3 | 0 | 3 |  | HOMOCYSDEGR-PWY | 4 | 4 | 4 |
| PWY-5458 | 3 | 0 | 2 |  | DETOX1-PWY-1 | 4 | 4 | 4 |
| PWY-5315 | 3 | 0 | 2 |  | PWY-7409 | 4 | 4 | 4 |
| SULFUROX-PWY | 3 | 0 | 2 |  | ARG-PRO-PWY | 4 | 4 | 4 |
| PWY-5515 | 3 | 0 | 2 |  | PWY-5659 | 4 | 4 | 4 |
| PWY-5080 | 4 | 0 | 4 |  | PWY-3781 | 4 | 4 | 4 |
| MGLDLCTANA-PWY | 4 | 0 | 3 |  | PWY-7218 | 4 | 4 | 4 |
| PWY-6117 | 5 | 0 | 3 |  | UDPNACETYLGALSYN-PWY | 4 | 4 | 4 |
| PWY-6369 | 9 | 1 | 9 |  | PROSYN-PWY | 4 | 4 | 4 |
| PWY-6368 | 9 | 1 | 9 |  | PWY-2261 | 4 | 3 | 3 |
| PWY-4361 | 7 | 1 | 5 |  | PWY-5059 | 4 | 3 | 3 |
| PWY-6261 | 15 | 2 | 10 |  | HEMESYN2-PWY | 4 | 3 | 3 |
| PWY-5285 | 3 | 1 | 3 |  | PWY1F-823 | 4 | 3 | 3 |
| PWY-6364 | 3 | 1 | 3 |  | PWY-621 | 4 | 3 | 3 |
| PROPIONMET-PWY | 3 | 1 | 3 |  | PWY490-3 | 4 | 3 | 3 |
| PWY-6307 | 4 | 1 | 3 |  | PWY-801 | 4 | 3 | 3 |
| DTDPRHAMSYN-PWY | 4 | 1 | 3 |  | PWY-6606 | 4 | 3 | 3 |
| PWY-5676 | 5 | 1 | 3 |  | PWY-5189 | 4 | 3 | 3 |
| PWY-401 | 5 | 2 | 5 |  | PWY-6305 | 4 | 3 | 3 |
| PWY-5329 | 2 | 1 | 2 |  | PWY-5384 | 4 | 3 | 3 |
| PWY-5996 | 2 | 1 | 2 |  | COA-PWY | 4 | 3 | 3 |
| PWY-5738 | 2 | 1 | 2 |  | PWY-7397 | 4 | 3 | 3 |
| PWY-4621 | 2 | 1 | 2 |  | PWY-5695 | 4 | 3 | 3 |
| GDPRHAMSYN-PWY | 2 | 1 | 2 |  | PWY-5750 | 4 | 3 | 3 |
| PWY-6613 | 2 | 1 | 2 |  | PWY-6689 | 5 | 5 | 5 |
| PWY-5760 | 2 | 1 | 2 |  | PWY-6124 | 5 | 5 | 5 |
| PWY-5054 | 3 | 1 | 2 |  | SALVADEHYPOX-PWY | 5 | 5 | 5 |
| PWY-4841 | 3 | 1 | 2 |  | PWY-3341 | 5 | 5 | 5 |
| PWY-6907 | 3 | 1 | 2 |  | PWY1F-FLAVSYN | 5 | 5 | 5 |
| PWY-6580 | 3 | 1 | 2 |  | PWY66-422 | 5 | 5 | 5 |
| PWY-5947 | 3 | 1 | 2 |  | NONOXIPENT-PWY | 5 | 5 | 5 |
| PWY66-368 | 3 | 1 | 2 |  | HOMOSER-METSYN-PWY | 5 | 5 | 5 |
| PWY-6181 | 3 | 1 | 2 |  | PWY-5905 | 5 | 5 | 5 |
| PWY-7206 | 3 | 1 | 2 |  | PWY-6317 | 5 | 5 | 5 |
| PWY-7375 | 3 | 1 | 2 |  | PLPSAL-PWY | 5 | 5 | 5 |
| PWY1-3 | 3 | 1 | 2 |  | PWY-2161 | 5 | 5 | 5 |
| PWY-3861 | 4 | 2 | 4 |  | PWY-7411 | 5 | 5 | 5 |
| PWY-6367 | 4 | 2 | 4 |  | PWY-7185 | 5 | 5 | 5 |
| DISSULFRED-PWY | 5 | 2 | 4 |  | GLUTORN-PWY | 5 | 5 | 5 |
| PWY3O-355 | 6 | 3 | 6 |  | PWY-6147 | 5 | 5 | 5 |
| PWY66-375 | 6 | 2 | 4 |  | PWY-6886 | 5 | 5 | 5 |
| PWY-5972 | 6 | 2 | 4 |  | PWY-6959 | 5 | 5 | 5 |
| PWY-3385 | 6 | 2 | 4 |  | PWY-6351 | 5 | 5 | 5 |
| PWY-5068 | 6 | 2 | 4 |  | PWY-4984 | 5 | 5 | 5 |
| PWY-7224 | 6 | 2 | 4 |  | PWY-5973 | 5 | 5 | 5 |
| P108-PWY | 7 | 2 | 4 |  | PWY-241 | 5 | 4 | 4 |
| PWY-6121 | 5 | 3 | 5 |  | PWY-6936 | 5 | 4 | 4 |
| FASYN-ELONG-PWY | 5 | 3 | 5 |  | PWY-6598 | 5 | 4 | 4 |
| GLYCOCAT-PWY | 8 | 3 | 5 |  | PWY-6554 | 5 | 4 | 4 |
| PWY-7654 | 11 | 5 | 8 |  | COA-PWY-1 | 5 | 4 | 4 |
| PWY-43 | 3 | 2 | 3 |  | PWY-3801 | 5 | 4 | 4 |
| PWY66-21 | 3 | 2 | 3 |  | PWY-7539 | 5 | 4 | 4 |
| ALACAT2-PWY | 3 | 2 | 3 |  | GLUCOSE1PMETAB-PWY | 5 | 3 | 3 |
| ARGDEG-III-PWY | 3 | 2 | 3 |  | TYRFUMCAT-PWY | 5 | 3 | 3 |
| PWY-3821 | 3 | 2 | 3 |  | PWY-5177 | 5 | 3 | 3 |
| SERDEG-PWY | 3 | 2 | 3 |  | PWY-7111 | 5 | 3 | 3 |
| ASPARAGINE-DEG1-PWY-1 | 3 | 2 | 3 |  | PWY-6992 | 5 | 3 | 3 |
| PWY0-1535 | 3 | 2 | 3 |  | UDPNAGSYN-PWY | 5 | 3 | 3 |
| PWY-5668 | 3 | 2 | 3 |  | LEU-DEG2-PWY | 6 | 6 | 6 |
| NAD-BIOSYNTHESIS-II | 3 | 2 | 3 |  | LEUSYN-PWY | 6 | 6 | 6 |
| PWY-3462 | 3 | 2 | 3 |  | PWY-7003 | 6 | 6 | 6 |
| METHIONINE-DEG1-PWY | 3 | 2 | 3 |  | GLYOXYLATE-BYPASS | 6 | 6 | 6 |
| PWY-6608 | 4 | 2 | 3 |  | PWY-5188 | 6 | 6 | 6 |
| PWY-7761 | 4 | 2 | 3 |  | PYRIDNUCSYN-PWY | 6 | 6 | 6 |
| PWY-7288 | 5 | 2 | 3 |  | PWY-5686 | 6 | 6 | 6 |
| P221-PWY | 5 | 2 | 3 |  | PWY0-862 | 6 | 6 | 6 |
| PWY6666-2 | 5 | 2 | 3 |  | PWY-5989 | 6 | 6 | 6 |
| PWY-7216 | 5 | 2 | 3 |  | ILEUDEG-PWY | 6 | 6 | 6 |
| PWY-6583 | 8 | 4 | 6 |  | PWY-6163 | 6 | 6 | 6 |
| PWY-735 | 19 | 10 | 15 |  | TRPSYN-PWY | 6 | 6 | 6 |
| PWY-5994 | 31 | 20 | 30 |  | PWY-7727 | 6 | 5 | 5 |
| PWY-7384 | 10 | 5 | 7 |  | PWY-7725 | 6 | 5 | 5 |
| PWY66-389 | 4 | 3 | 4 |  | PWY-6370 | 6 | 4 | 4 |
| PWY-5653 | 4 | 3 | 4 |  | PWY-5109 | 6 | 4 | 4 |
| PWY-5675 | 4 | 3 | 4 |  | PWY-4981 | 6 | 4 | 4 |
| PWY-7663 | 4 | 3 | 4 |  | RUMP-PWY | 6 | 4 | 4 |
| PWY-5041 | 4 | 3 | 4 |  | PWY-6123 | 6 | 4 | 4 |
| ARGININE-SYN4-PWY | 4 | 3 | 4 |  | PWY-1422 | 7 | 7 | 7 |
| PWY-6113 | 4 | 3 | 4 |  | PWY-922 | 7 | 7 | 7 |
| PWY0-1264 | 4 | 3 | 4 |  | TRIGLSYN-PWY | 7 | 7 | 7 |
| PWY-6122 | 5 | 3 | 4 |  | PWY-5097 | 7 | 7 | 7 |
| PWY-6277 | 5 | 3 | 4 |  | PWY-7426 | 7 | 7 | 7 |
| PWY66-367 | 5 | 3 | 4 |  | PWY-6823 | 7 | 7 | 7 |
| PWY-6435 | 5 | 3 | 4 |  | ILEUSYN-PWY | 7 | 7 | 7 |
| PWY-5437 | 6 | 3 | 4 |  | PWY-7198 | 7 | 6 | 6 |
| PWY-7118 | 6 | 3 | 4 |  | PWY-2942 | 7 | 6 | 6 |
| PWY-6883 | 6 | 3 | 4 |  | PWY-6922 | 7 | 6 | 6 |
| PYRIDNUCSAL-PWY | 6 | 3 | 4 |  | PWY-7053 | 7 | 5 | 5 |
| PWY-6854 | 7 | 3 | 4 |  | PWY-6871 | 7 | 5 | 5 |
| PWY-6313 | 7 | 3 | 4 |  | SUCSYN-PWY | 7 | 5 | 5 |
| PWY-3661 | 7 | 3 | 4 |  | FAO-PWY | 7 | 5 | 5 |
| PWY-7664 | 14 | 10 | 13 |  | PWY-6174 | 7 | 5 | 5 |
| PWY-7269 | 5 | 4 | 5 |  | PWY-7601 | 7 | 4 | 4 |
| PWY-6362 | 5 | 4 | 5 |  | PWY-7383 | 7 | 4 | 4 |
| PWY-5136 | 5 | 4 | 5 |  | PWY-5103 | 7 | 4 | 4 |
| PWY-6531 | 5 | 4 | 5 |  | CENTFERM-PWY | 7 | 4 | 4 |
| PWY-7268 | 5 | 4 | 5 |  | VALDEG-PWY | 8 | 8 | 8 |
| PWY-5083 | 6 | 4 | 5 |  | CITRULBIO-PWY | 8 | 8 | 8 |
| PWY-7094 | 6 | 4 | 5 |  | PWY-7210 | 8 | 8 | 8 |
| PWY66-391 | 7 | 4 | 5 |  | PWY-6352 | 8 | 8 | 8 |
| PWY-7238 | 8 | 4 | 5 |  | PWY-6901 | 8 | 7 | 7 |
| PWY-1042 | 10 | 8 | 10 |  | PWY-7124 | 8 | 7 | 7 |
| FERMENTATION-PWY | 16 | 8 | 10 |  | PWY-7391 | 8 | 7 | 7 |
| PWY-6527 | 7 | 5 | 6 |  | PWY-6596 | 8 | 6 | 6 |
| PWY-5514 | 7 | 5 | 6 |  | 1CMET2-PWY | 9 | 9 | 9 |
| PWY-7282 | 9 | 5 | 6 |  | PWY-7388 | 9 | 9 | 9 |
| PWY-7204 | 9 | 5 | 6 |  | ARGSYNBSUB-PWY | 9 | 9 | 9 |
| PWY-5381 | 11 | 5 | 6 |  | PWY-7115 | 9 | 9 | 9 |
| PWY-6728 | 18 | 11 | 13 |  | PWY-6282 | 9 | 9 | 9 |
| PWY-6519 | 11 | 7 | 8 |  | PWY-7184 | 9 | 9 | 9 |
| P105-PWY | 11 | 9 | 10 |  | PWY-6545 | 9 | 8 | 8 |
| PWY66-399 | 12 | 10 | 11 |  | PWY-6549 | 9 | 8 | 8 |
| P122-PWY | 18 | 13 | 14 |  | PWY-5154 | 9 | 7 | 7 |
| PWY-5971 | 31 | 28 | 30 |  | PWY-7400 | 9 | 7 | 7 |
| PWY-6964 | 2 | 2 | 2 |  | PWY-1861 | 9 | 7 | 7 |
| GLYCOLYSIS-E-D | 2 | 2 | 2 |  | DAPLYSINESYN-PWY | 9 | 6 | 6 |
| DETOX1-PWY | 2 | 2 | 2 |  | PWY-2941 | 9 | 6 | 6 |
| PWY-2301 | 2 | 2 | 2 |  | P341-PWY | 9 | 6 | 6 |
| PWY-6146 | 2 | 2 | 2 |  | PWY-6168 | 9 | 6 | 6 |
| PWYQT-4427 | 2 | 2 | 2 |  | REDCITCYC | 9 | 5 | 5 |
| GLUT-REDOX-PWY | 2 | 2 | 2 |  | CODH-PWY | 9 | 5 | 5 |
| PWY-7343 | 2 | 2 | 2 |  | PWY-7385 | 9 | 5 | 5 |
| MALATE-ASPARTATE-SHUTTLE-PWY | 2 | 2 | 2 |  | ANAGLYCOLYSIS-PWY | 10 | 10 | 10 |
| 2PHENDEG-PWY | 2 | 2 | 2 |  | PWY-7250 | 10 | 10 | 10 |
| PWY-5943 | 2 | 2 | 2 |  | PWY-7434 | 10 | 10 | 10 |
| PWY-4261 | 2 | 2 | 2 |  | HISTSYN-PWY | 10 | 10 | 10 |
| PWY-6543 | 2 | 2 | 2 |  | PWY-7117 | 10 | 9 | 9 |
| PWY-7226 | 2 | 2 | 2 |  | PWY-6787 | 10 | 6 | 6 |
| PWY-6118 | 2 | 2 | 2 |  | PWY-5484 | 11 | 11 | 11 |
| PWY-5340 | 2 | 2 | 2 |  | PWY-5913 | 11 | 10 | 10 |
| PWY-7227 | 2 | 2 | 2 |  | PWY-3841 | 11 | 10 | 10 |
| PWY-6019 | 2 | 2 | 2 |  | PWY66-398 | 11 | 10 | 10 |
| HSERMETANA-PWY | 2 | 2 | 2 |  | PWY-2201 | 12 | 12 | 12 |
| PWY-7417 | 2 | 2 | 2 |  | GLYCOLYSIS | 12 | 12 | 12 |
| PWY66-423 | 2 | 2 | 2 |  | P185-PWY | 12 | 11 | 11 |
| PWY-6952 | 2 | 2 | 2 |  | PWY-6969 | 12 | 10 | 10 |
| PWY-5921 | 2 | 2 | 2 |  | GLUCONEO-PWY | 13 | 13 | 13 |
| PWY-7205 | 2 | 2 | 2 |  | PWY-7606 | 14 | 14 | 14 |
| PWY-3981 | 2 | 2 | 2 |  | MANNOSYL-CHITO-DOLICHOL-BIOSYNTHESIS | 19 | 19 | 19 |
| PWY-5537 | 2 | 2 | 2 |  | TRNA-CHARGING-PWY | 21 | 21 | 21 |
| PWY-6001 | 2 | 2 | 2 |  | CALVIN-PWY | 13 | 13 | 12 |
| PWY-66 | 2 | 2 | 2 |  | P124-PWY | 15 | 12 | 11 |
| PWY0-1466 | 2 | 2 | 2 |  | PWY-5723 | 10 | 10 | 9 |
| THIOREDOX-PWY | 2 | 2 | 2 |  | PWY-6142 | 12 | 10 | 9 |
| PWY4FS-8 | 2 | 2 | 2 |  | P23-PWY | 12 | 10 | 9 |
| BETSYN-PWY | 2 | 2 | 2 |  | PWY-7560 | 9 | 9 | 8 |
| PWY-5670 | 2 | 2 | 2 |  | TCA | 10 | 9 | 8 |
| PWY66-366 | 2 | 2 | 2 |  | NONMEVIPP-PWY | 9 | 8 | 7 |
| PWY-6012 | 2 | 2 | 2 |  | RIBOSYN2-PWY | 9 | 8 | 7 |
| PWY-7494 | 2 | 2 | 2 |  | PWY-5690 | 9 | 8 | 7 |
| PWYQT-4429 | 2 | 2 | 2 |  | PWY-882 | 8 | 7 | 6 |
| FASYN-INITIAL-PWY | 2 | 2 | 2 |  | PWY-7254 | 9 | 7 | 6 |
| GLUTATHIONESYN-PWY | 2 | 2 | 2 |  | PWY-6863 | 11 | 7 | 6 |
| CHOLINE-BETAINE-ANA-PWY | 2 | 2 | 2 |  | PWY-702 | 6 | 6 | 5 |
| PWY-6 | 2 | 2 | 2 |  | CHLOROPHYLL-SYN | 9 | 6 | 5 |
| HOMOSER-THRESYN-PWY | 2 | 2 | 2 |  | PWY-181 | 9 | 6 | 5 |
| PWY-6164 | 2 | 2 | 2 |  | PWY0-166 | 13 | 12 | 10 |
| PWY0-1182 | 2 | 2 | 2 |  | P42-PWY | 7 | 5 | 4 |
| CYSTSYN-PWY | 2 | 2 | 2 |  | PWY-2221 | 9 | 5 | 4 |
| PWY-5278 | 2 | 2 | 2 |  | PWY-7159 | 9 | 5 | 4 |
| PWY-5697 | 2 | 2 | 2 |  | PWY-4041 | 4 | 4 | 3 |
| PWY-6963 | 2 | 2 | 2 |  | PWY-7220 | 4 | 4 | 3 |
| BSUBPOLYAMSYN-PWY | 2 | 2 | 2 |  | PWY-7222 | 4 | 4 | 3 |
| TRESYN-PWY | 2 | 2 | 2 |  | PANTO-PWY | 4 | 4 | 3 |
| PWY-5966 | 2 | 2 | 2 |  | PWY-7432 | 4 | 4 | 3 |
| PWY4FS-7 | 2 | 2 | 2 |  | PWY-6398 | 5 | 4 | 3 |
| PWY-5046 | 3 | 3 | 3 |  | PWY-6902 | 5 | 4 | 3 |
| P21-PWY | 3 | 3 | 3 |  | PWY66-387 | 6 | 4 | 3 |
| LCYSDEG-PWY | 3 | 3 | 3 |  | PWY66-388 | 7 | 4 | 3 |
| HOMOSERSYN-PWY | 3 | 3 | 3 |  | PWY-7356 | 7 | 4 | 3 |
| PWY-5084 | 3 | 3 | 3 |  | PWY-5123 | 3 | 3 | 2 |
| PWY-4081 | 3 | 3 | 3 |  | PWY-4302 | 3 | 3 | 2 |
| PWY-4101 | 3 | 3 | 3 |  | PWY-6120 | 3 | 3 | 2 |
| TYRSYN | 3 | 3 | 3 |  | PWY-3461 | 3 | 3 | 2 |
| PWY-6363 | 3 | 3 | 3 |  | NADPHOS-DEPHOS-PWY | 3 | 3 | 2 |
| PWY-7176 | 3 | 3 | 3 |  | PWY-5 | 4 | 3 | 2 |
| PWY-1801 | 3 | 3 | 3 |  | PWY-6859 | 4 | 3 | 2 |
| ARGASEDEG-PWY | 3 | 3 | 3 |  | PWY-6609 | 4 | 3 | 2 |
| SERSYN-PWY | 3 | 3 | 3 |  | PWY-6803 | 4 | 3 | 2 |
| PYRUVDEHYD-PWY | 3 | 3 | 3 |  | PWY-5194 | 4 | 3 | 2 |
| PROUT-PWY | 3 | 3 | 3 |  | PWY-101 | 4 | 3 | 2 |
| PWY-7177 | 3 | 3 | 3 |  | PWY-5981 | 5 | 3 | 2 |
| PWY-6481 | 3 | 3 | 3 |  | PWY-7039 | 5 | 3 | 2 |
| PWY-7416 | 3 | 3 | 3 |  | PWY-7187 | 7 | 6 | 4 |
| PWY-1722 | 3 | 3 | 3 |  | PWY-6309 | 11 | 6 | 4 |
| GLYCLEAV-PWY | 3 | 3 | 3 |  | PWY66-428 | 2 | 2 | 1 |
| PHESYN | 3 | 3 | 3 |  | ARG-GLU-PWY | 2 | 2 | 1 |
| PWY-6632 | 3 | 3 | 3 |  | NADPHOS-DEPHOS-PWY-1 | 2 | 2 | 1 |
| PWY-7193 | 3 | 3 | 3 |  | PWY-5538 | 2 | 2 | 1 |
| PWY66-162 | 3 | 3 | 3 |  | CITRULLINE-DEG-PWY | 2 | 2 | 1 |
| PWY0-1507 | 3 | 3 | 3 |  | ARGSPECAT-PWY | 2 | 2 | 1 |
| PWY-5691 | 3 | 3 | 3 |  | TREHALOSESYN-PWY | 2 | 2 | 1 |
| OXIDATIVEPENT-PWY | 3 | 3 | 3 |  | PWY-881 | 2 | 2 | 1 |
| PWY-6614 | 3 | 3 | 3 |  | SO4ASSIM-PWY | 2 | 2 | 1 |
| PWY-381 | 3 | 3 | 3 |  | PWY-6000 | 2 | 2 | 1 |
| PWY-6683 | 3 | 2 | 2 |  | PWY-6897 | 3 | 2 | 1 |
| SULFMETII-PWY | 3 | 2 | 2 |  | RIBOKIN-PWY | 3 | 2 | 1 |
| PWY0-1545 | 3 | 2 | 2 |  | PWY-6983 | 3 | 2 | 1 |
| ANAPHENOXI-PWY | 3 | 2 | 2 |  | PWY490-4 | 3 | 2 | 1 |
| PWY-5938 | 3 | 2 | 2 |  | PWY-6610 | 3 | 2 | 1 |
| PWY-5386 | 3 | 2 | 2 |  | PHOSLIPSYN2-PWY | 3 | 2 | 1 |
| PWY-6073 | 3 | 2 | 2 |  | PWY-6788 | 3 | 2 | 1 |
| PWY-6917 | 3 | 2 | 2 |  | PWY0-901 | 3 | 2 | 1 |
| PWY-6908 | 3 | 2 | 2 |  | ADENOSYLHOMOCYSCAT-PWY | 3 | 2 | 1 |
| PWY-7618 | 3 | 2 | 2 |  | PWY0-1319 | 4 | 4 | 2 |
| PWY-701 | 3 | 2 | 2 |  | PWY-7726 | 13 | 13 | 6 |
| PWY-5108 | 3 | 2 | 2 |  | PWY-5663 | 3 | 3 | 1 |
| PWY-3881 | 3 | 2 | 2 |  | PWY3O-450 | 3 | 3 | 1 |
| PWY-3641 | 3 | 2 | 2 |  | PWY-3561 | 3 | 3 | 1 |
| PWY-5269 | 3 | 2 | 2 |  | PWY-2181 | 4 | 3 | 1 |
| ETHYL-PWY | 3 | 2 | 2 |  | PWY-5064 | 5 | 3 | 1 |
| PWY66-161 | 3 | 2 | 2 |  | LIPAS-PWY | 4 | 4 | 1 |
| PWY-6389 | 3 | 2 | 2 |  | PWY-6281 | 4 | 4 | 1 |
| PWY-5390 | 3 | 2 | 2 |  | PWY-7728 | 5 | 4 | 1 |
| PWY-7394 | 3 | 2 | 2 |  | PWY3DJ-11470 | 5 | 4 | 1 |
| PWY1F-353 | 3 | 2 | 2 |  | LIPASYN-PWY | 5 | 4 | 1 |
| PWY-6416 | 3 | 2 | 2 |  | LYSINE-DEG1-PWY | 5 | 5 | 1 |
| TRYPDEG-PWY | 3 | 2 | 2 |  | PWY-7511 | 9 | 9 | 1 |
| PWY-2723 | 3 | 2 | 2 |  | PWY-6599 | 2 | 2 | 0 |
| PWY-4983 | 3 | 2 | 2 |  | PWY0-1587 | 2 | 2 | 0 |
| CYSTEINE-DEG-PWY | 3 | 2 | 2 |  | ENTNER-DOUDOROFF-PWY | 2 | 2 | 0 |
| CYANCAT-PWY | 3 | 2 | 2 |  | GLUDEG-II-PWY | 2 | 2 | 0 |
| PWY-4381 | 3 | 2 | 2 |  | PWY-6825 | 3 | 3 | 0 |
| PWY-7586 | 3 | 2 | 2 |  | ARGDEG-V-PWY | 3 | 3 | 0 |
| PWY-7380 | 3 | 2 | 2 |  | PWY-5063 | 3 | 3 | 0 |
| PWY-5661 | 3 | 2 | 2 |  | PWY-5901 | 3 | 2 | 0 |
| HEME-BIOSYNTHESIS-II | 4 | 4 | 4 |  | PWY-7587 | 3 | 2 | 0 |
| PWY-7219 | 4 | 4 | 4 |  | PWY-7092 | 3 | 2 | 0 |
| PWY-561 | 4 | 4 | 4 |  | PWY-6927 | 5 | 3 | 0 |
| PWY-7221 | 4 | 4 | 4 |  |  |  |  |  |

### Supporting Information References
